# Reenacting a mouse genetic evolutionary arms race in yeast reveals SLXL1/SLX compete with SLY1/2 for binding to Spindlins

**DOI:** 10.1101/2024.10.18.619120

**Authors:** Martin F. Arlt, Alyssa N. Kruger, Callie M. Swanepoel, Jacob L. Mueller

## Abstract

The house mouse X and Y chromosomes have recently acquired high copy number, rapidly evolving gene families representing an evolutionary arms race. This arms race between proteins encoded by X-linked *Slxl1*/*Slx* and Y-linked *Sly* gene families can distort male offspring sex ratio, but how these proteins compete remains unknown. Here, we report how *Slxl1*/*Slx* and *Sly* encoded proteins compete in a protein family-specific and dose-dependent manner using yeast. Specifically, SLXL1 competes with SLY1 and SLY2 for binding to the Spindlin SPIN1. Similarly, SLX competes with SLY2 for binding the Spindlin SSTY2. These competitions are driven by the N-termini of SLXL1, SLX, SLY1, and SLY2 binding to the third Tudor domains of SPIN1 and SSTY2. SLY1 and SLY2 form homo- and heterodimers, suggesting the competition is between complex multimers. Residues under positive selection mapping to the interaction domains and rapid exon gain/loss are consistent with competition between the X- and Y-linked gene families. Our findings support a model in which dose-dependent competition of these X- and Y-linked encoded proteins to bind Spindlins occurs in haploid X- and Y-spermatids to influence X-versus Y-sperm fitness and thus sex ratio.

**Significance Statement:** In house mouse, an evolutionary arms race between proteins encoded by the X-linked *Slxl1/Slx* and Y-linked *Sly* gene families during spermatogenesis can distort male offspring sex ratio, but how these proteins compete remains unknown. We report how SLXL1/SLX competes with SLY1/SLY2 by demonstrating their dose-dependent competitive binding to Spindlins, the key protein domains and rapidly evolving residues and exons that drive the competition, and how the competition is likely between complex multimers. Our findings have broad implications for the mechanics of evolutionary arms and how competition between sex chromosomes influences X-versus Y-sperm fitness and sex ratio.

## Introduction

In evolutionary arms races, co-evolving genes develop adaptations and counter-adaptations against each other. These arms races can evolve in various ways, including between species, such as in host-pathogen interactions, or between chromosomes within a species, such as the X and Y chromosome (1–3). The nature of competition in arms races and how they drive proteins’ rapid evolution remains unclear. In the house mouse, competition between X-chromosome and Y-chromosome-bearing sperm is driven by an evolutionary arms race between the X-linked *Slxl1*/*Slx* and the Y-linked *Sly* gene families (4–9). These gene families arose from autosomal *Synaptonemal complex protein-3* (*Sycp3*) (6, 10), collectively called *<u>S</u>ycp3*-like genes on the <u>X</u> and <u>Y</u> chromosomes. *Slxl1*/*Slx* and *Sly* are present in tens to hundreds of gene copies and expressed exclusively in haploid X and Y-bearing spermatids (5, 6, 8, 11, 12). The primary evidence that *Slxl1*/*Slx* and *Sly* compete in an evolutionary arms race is that loss of function studies of the X- or Y-linked copies results in more male or female offspring, respectively (termed sex ratio distortion) (4, 5, 8). This sex ratio distortion is thought to be a consequence of X-versus Y-spermatid gene expression differences that alter the fitness of X-versus Y-sperm to fertilize an egg (4, 5, 7–9, 12). Our laboratory identified the top three candidate protein interactors of SLXL1/SLX and SLY1/2 as the known and putative Spindlin chromatin readers, SPIN1, SSTY1, and SSTY2 (8). This finding led to our speculation that SLXL1/SLX and SLY1/2 compete for Spindlin binding to influence X-versus Y-spermatid gene expression, leading to X- and Y-sperm fitness and thus sex ratio distortion (8). However, it remains unclear whether the SYCP3-like protein interactions are direct or indirect, protein-family-specific (i.e., SLX vs. SLXL1 or SLY1 vs. SLY2), or whether the X- and Y-encoded proteins interact with the same or different substrates (i.e. SLXL1 vs. SLY1).

We hypothesize two main adaptations/counter-adaptations driving the *Slxl1/Slx* versus *Sly* evolutionary arms race. First, sequence changes have created an optimal set of protein interactions for each gene family. Second, the high gene copy number reflects the importance of gene dosage in influencing competition. This study tests our hypothesis via three critical aspects of the SLXL1/SLX versus SLY1/2 arms race using *in vitro* and computational approaches. First, we identify these proteins’ binding substrates and interaction domains using yeast two-hybrid (Y2H). Next, we demonstrate dose-dependent competition for protein interactions using a modified Y3H system to establish the molecular mechanism underlying the competition. Finally, we investigate the molecular evolution of *Slx*, *Slxl1*, and *Sly* to find how the acquisition, protein sequence changes, and gene amplification during the arms race are related to the protein interactions. These observations provide critical mechanistic insight into how competition between SLXL1/SLX and SLY1/2 influences sperm fitness via an X-versus Y-chromosome evolutionary arms race during mouse spermatogenesis. Exploring how this arms race evolves and competes provides insights into innovative molecular strategies, driven by strong positive Darwinian selection, influencing fertility and chromosome evolution.

## Results

### SYCP3-like proteins are predicted to compete for binding to Spindlins

Our previous immunoprecipitation of SLX and SLXL1 from mouse testis followed by mass spectrometry to identify Spindlins SPIN1 and SSTY1/2 as candidate binding proteins (8). SLY1 also interacts with SPIN1 and SSTY1/2 (13), as well as other candidate-interacting proteins (9, 12). The observation that the SYCP3-like proteins all appear to interact with SPIN1 and SSTY1/2 makes the Spindlins an attractive target for competition. It is unknown whether SLX, SLXL1, and SLY1/2 interactions with Spindlins are direct and whether SLX and SLXL1 interact with the same proteins as the related Y-linked proteins, SLY1 and SLY2.

We assessed the binding of SLXL1, SLX, SLY1, and SLY2 with candidate proteins, using AlphaFold Multimer (14) to test all pairwise interactions (Fig. 1a). Consistent with previous studies (8, 9, 13), each of the SYCP3-like proteins is predicted to bind SPIN1 and SSTY1/2 (Fig. 1b, Supp. Fig. 1a). (8, 13, 15–18) SLXL1, SLX, SLY1, and SLY2 have similar structures with two predominant features: a pair of β-strands near the N-terminus and a long C-terminal α-helix. SPIN1 and SSTY1/2 are made up of three Tudor-like domains (Tudor1, Tudor2, Tudor3) (17, 19) (Fig. 1c, Supp. Fig. 1b). Based on the prediction that all four SYCP3-like proteins interact with these three Spindlins, we hypothesize the X-linked SLXL1 and SLX compete with the Y-linked counterparts SLY1 and SLY2 for binding to SPIN1/SSTY1/SSTY2.

**Figure 1.**
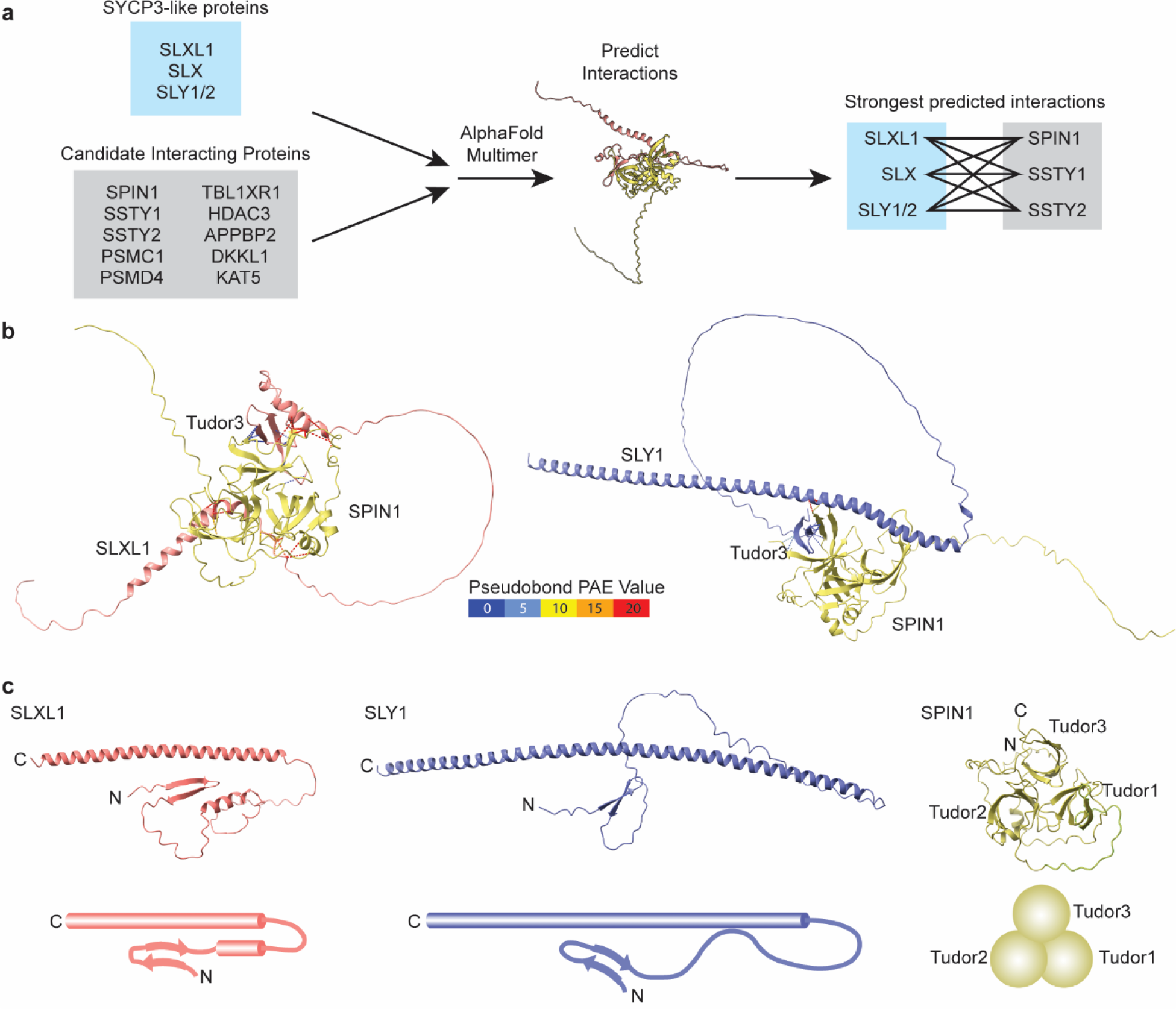
SYCP3-like proteins are predicted to bind Spindlins. (a) Candidate SYCP3-like protein interactors from previous studies (8, 9, 12) tested for predicted pairwise interactions using AlphaFold Multimer (14). The most robust predicted direct interactions are between SYCP3-like and Spindlin proteins. (b) An example of an AlphaFold Multimer predicted SPIN1 binding to SLXL1 (left) and SLY1 (right), with (c) associated 3D structure diagrams.

### SLXL1/SLY1/SLY2 bind to SPIN1, and SLX/SLY2 bind to SSTY2

We first asked if these proteins directly interact in a Y2H assay to test the hypothesis that SLXL1 and SLX compete with SLY1/2 for binding to Spindlins. In contrast to AlphaFold Multimer predictions, we find SPIN1 interacts with SLXL1, SLY1, and SLY2, but not with SLX. SSTY2 interacts with SLX and SLY2 but not with SLXL1 or SLY1. SSTY1 does not interact with SYCP3-like proteins (Fig. 2a, b). Y2H with the Gal4 activation domain (AD) and binding domain (BD) swapped recapitulated SPIN1 interactions with SLY1/2 and SSTY2 interactions with SLY2 (Supp. Fig. 3). However, fusing SLXL1/SLX to a BD results in autoactivation of reporter genes, preventing us from repeating the Y2H experiments with SLXL1/SLX in the bait plasmid (Supp. Fig. 2).

**Figure 2.**
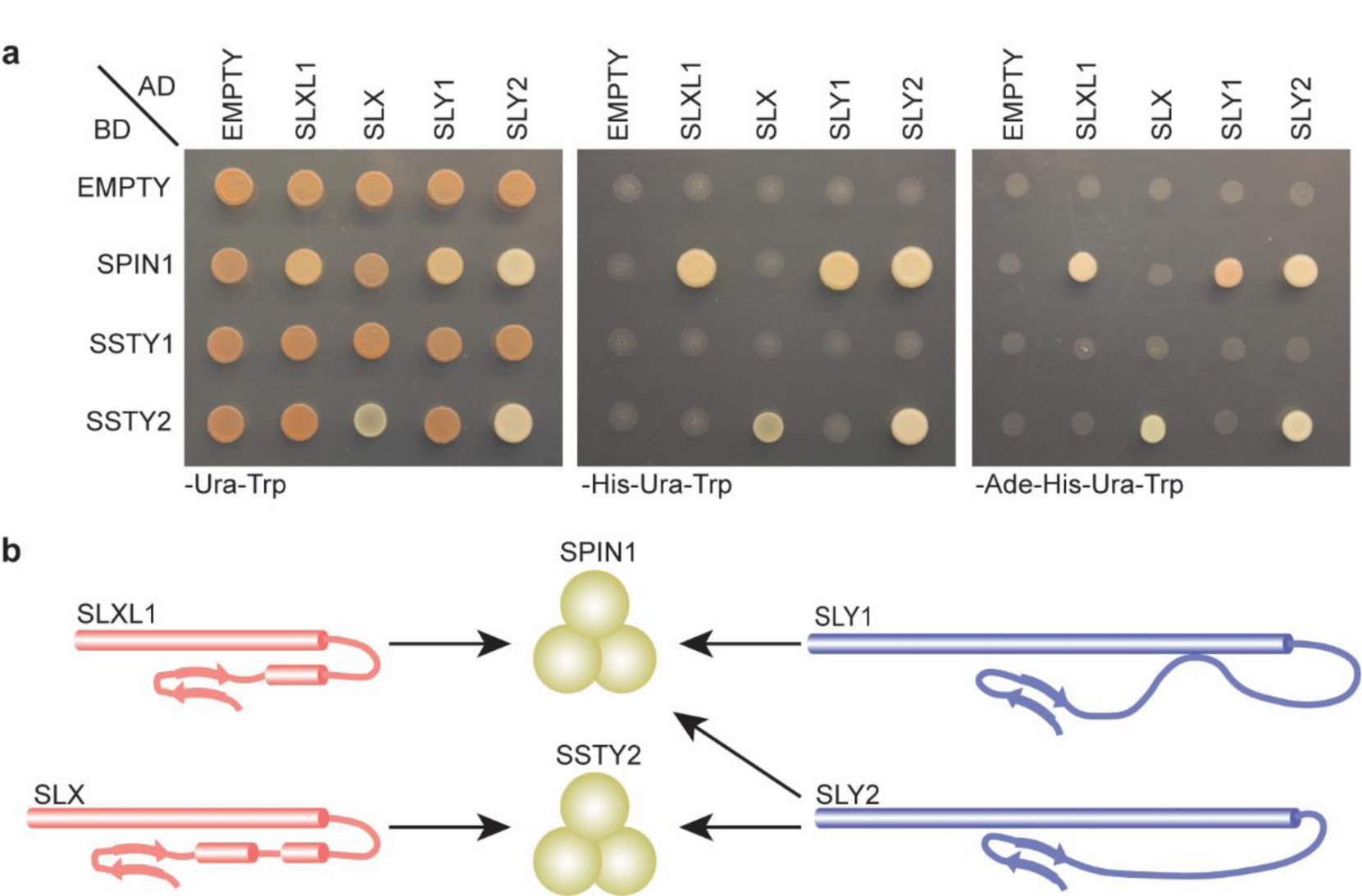
SLXL1/SLY1/SLY2 bind to SPIN1 and SLX/SLY2 bind to SSTY2. (a) SYCP3-like proteins tested for binding to Spindlins in Y2H spot assays. Growth on -Ura-Trp indicates the presence of BD and AD vectors. Growth on -His-Ura-Trp and -Ade-His-Ura-Trp indicates binding between BD- and AD-linked proteins. Negative controls: Empty vectors. (b) Summary of Y2H binding results (arrows).

To evaluate whether SLXL1/SLX/SLY1/SLY2 generally bind to Spindlins, we tested for interactions with four additional Spindlins (SSTX, SPIN2C, SPIN4, and GM4868). While AlphaFold Multimer predicts pairwise interactions between SLXL1/SLX/SLY1/SLY2 and each of these Spindlins, no interactions were observed in our Y2H assay (Supp. Fig. 4). These data suggest SLXL1/SLY1/SLY2 interact specifically with SPIN1, SLX/SLY2 interact specifically with SSTY2, and SYCP3-like proteins do not interact with other Spindlins.

### N-terminal β-strands of SLXL1/SLX/SLY1/SLY2 are necessary and sufficient for binding to SPIN1 and SSTY2

With knowledge of the Spindlin binding partners of SLXL1, SLX, and SLY1/2, we can dissect what domains are essential for binding, which informs how they compete. The N-terminal β-strands of SLXL1, SLX, and SLY1/2 are predicted to bind to the Tudor3 domain of their partner Spindlin(s), suggesting that the interaction of these domains is the driver of competition (Fig. 3a, Supp. Fig. 5a). We tested if the N-terminal β-strands of SYCP3-like proteins are necessary for their interaction with SPIN1/SSTY2 using Y2H bait plasmids lacking the β-strands at amino acids 2-20 of SLXL1/SLX and 2-23 of SLY1/2 (Fig. 3b). We find that SYCP3-like proteins lacking this β-strand domain cannot bind to SPIN1 or SSTY2 (Fig. 3c).

**Figure 3.**
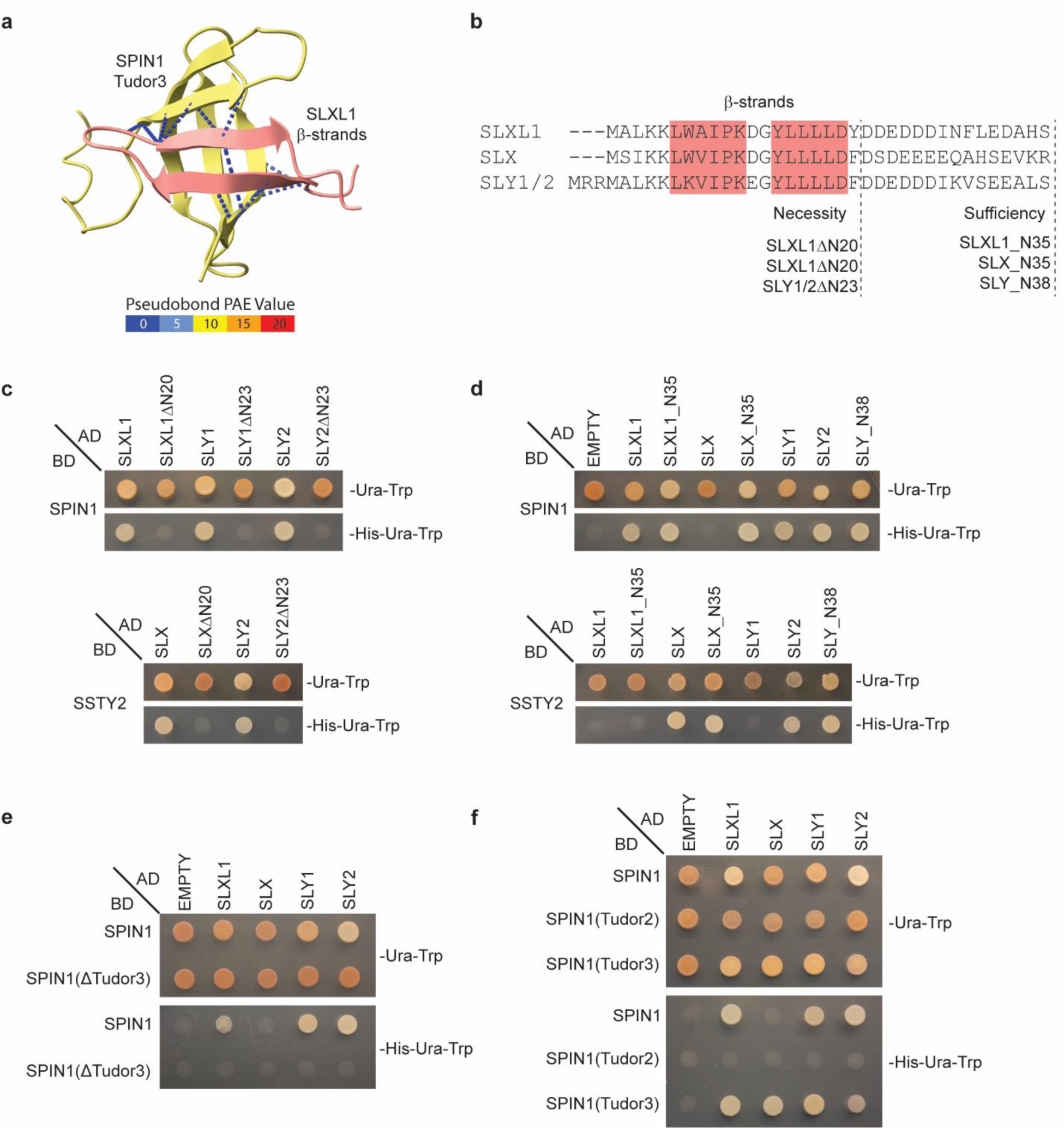
N-terminal β-strands of SYCP3-like proteins are necessary and sufficient for binding to SPIN1/SSTY2 and SPIN1 Tudor3 domain is necessary and sufficient for binding to SYCP3-like proteins. (a) AlphaFold Multimer predicted interaction between SLXL1 and SPIN1 (left) with a close-up of SLXL1 N-terminal β-strands and SPIN1 Tudor3. Dashed lines represent interresidue distances ≤ 3 Å. Intermolecular pseudobonds are color-coded by predicted aligned error (PAE) value as indicated. (b) N-terminal 35/38 amino acid alignment of SYCP3-like proteins with pink boxes highlighting β-strands. Dashed lines indicate amino acids deleted or included to test necessity and sufficiency, respectively, for binding in the Y2H assay. (c) Y2H spot assays demonstrate the N-terminal 20-23 amino acids of SYCP3-like proteins are necessary to interact with Spindlins. (d) Y2H spot assays demonstrate the N-terminal 35-38 amino acids of SYCP3-like proteins are sufficient to interact with SPIN1. (e) Y2H spot assay demonstrates that the SPIN1 Tudor3 domain is necessary for interacting with SYCP3-like proteins. (f) Y2H spot assay demonstrates SPIN1 Tudor3 domain alone is sufficient to interact with SYCP3-like proteins, while SPIN1 Tudor2 domain is not.

We then asked whether these N-terminal structures are sufficient for binding to SPIN1/SSTY2. We tested the ability of amino acids 1-35 of SLXL1 and SLX or amino acids 1-38 of SLY (which are identical between SLY1 and SLY2 isoforms) (Fig. 3b) to bind SPIN1 and SSTY2. We find the N-terminal β-strands of SLXL1/SLY and SLX/SLY are sufficient for binding to SPIN1 and SSTY2, respectively (Fig. 3d).

We also find that the N-terminal β-strands of SLX can interact with SPIN1, even though full-length SLX does not (Fig. 3d). This observation suggests that while SPIN1 can bind to the N-terminus of SLXL1, SLX, and SLY, other regions of these proteins contribute to the specificity of this interaction. In contrast, like full-length SLXL1, the SLXL1 N-terminus alone cannot interact with SSTY2, indicating specificity for that interaction is within the N-terminal 35 amino acids of SLXL1 (Fig. 3d).

### SPINDLIN Tudor3 domains are necessary and sufficient for binding to SYCP3-like proteins

With confirmation that the N-terminal domains of SYCP3-like proteins are necessary and sufficient for Spindlin interactions, we performed a reciprocal test: are the Spindlin Tudor3 domains necessary and sufficient for interacting with SYCP3-like proteins (Fig. 3a, Supp. Fig. 5a). SPIN1 lacking Tudor3 (SPIN1ΔTudor3) does not interact with full-length SLXL1/SLX/SLY1/SLY2 (Fig. 3e), indicating the SPIN1 Tudor3 domain is necessary for interactions with the SYCP3-like proteins. Similarly, SSTY2 with a deletion of Tudor3 (SSTY2ΔTudor3) does not interact with full-length SLX/SLY1/SLY2 (Supp. Fig. 5b). However, SSTY2ΔTudor3 does interact with SLXL1, even though full-length SSTY2 does not (Supp. Fig. 5b). We find that the interaction between SSTY2ΔTudor3 is dependent on the N-terminal 20 amino acids of SLXL1, as deleting these residues abrogates the binding (Supp. Fig. 5c). AlphaFold Multimer predicts that interactions between the N-terminal β-sheet of SLXL1 and both Tudor domains 1 and 2 of SSTY2 mediate the binding of SSTY2ΔTudor3 with full-length SLXL1 (Supp. Fig. 5d).

Next, we examined if SPIN1 or SSTY2 Tudor3 domains are sufficient for binding full-length SLXL1/SLX/SLY1/SLY2. We also tested if SPIN1 Tudor3 is sufficient for binding to the N-termini of the SYCP3-like proteins (SLXL_N35/SLX_N35/SLY_N38). We found the Tudor3 domain of either SPIN1 or SSTY2 is sufficient to interact with full-length SLXL1/SLX/SLY1/SLY2 (Fig. 3f, Supp. Fig. 5e). In addition, SPIN1 Tudor3 can bind to the N-termini alone of SLXL1/SLX/SLY1/SLY2 (Supp. Fig. 6). As a negative control, we showed the SPIN1 Tudor2 domain, which AlphaFold Multimer does not predict to interact with these proteins, cannot interact with SLXL1/SLX/SLY1/SLY2 (Fig. 3f). The interaction of SPIN1 Tudor3 and SSTY2 Tudor3, but not full-length proteins, with SLX and SLXL1/SLY1, respectively, indicates that while these Tudor3 domains are both necessary and sufficient for binding to SYCP3-like proteins, other SPINDLIN domains contribute to the interaction specificity.

### SLY1 and SLY2 can form homo- and hetero-dimers

With an improved understanding of how SYCP3-like proteins specifically interact with Spindlins, we questioned whether they compete not as dimers, but as more complex multimers. The basis for this reasoning is that SYCP3 can form homo-tetramers mediated by its α-helix (20), so we asked if SYCP3-like proteins form homo- or heterodimers. SLXL1/SLX/SLY1/SLY2 are predicted to form homodimers and heterodimers via the α-helical domains (Fig. 4a), using AlphaFold Multimer.

**Figure 4.**
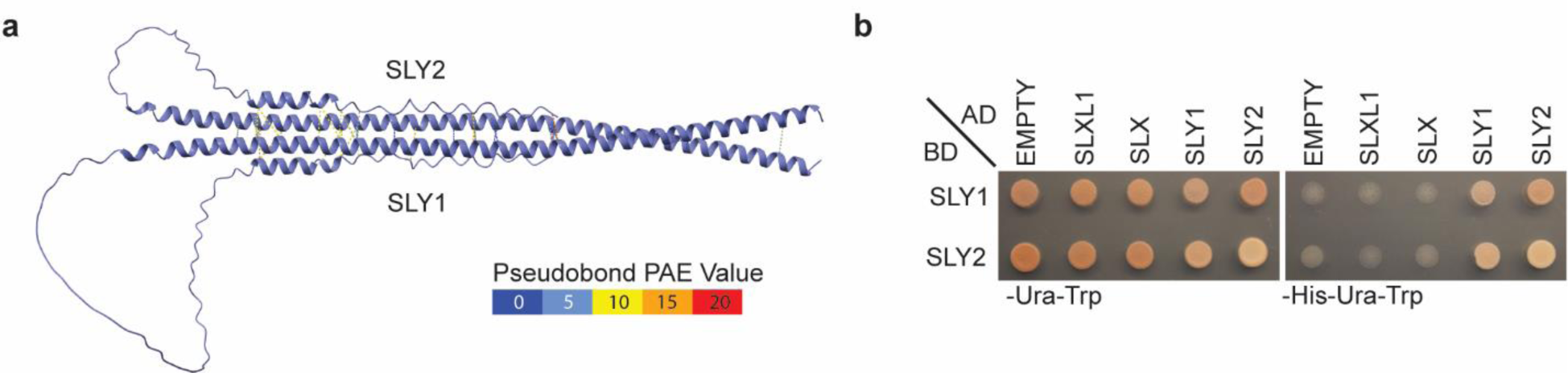
SLY1 and SLY2 can form homo- and hetero-dimers. (a) Predicted interactions between SLY1 and SLY2 based on AlphaFold Multimer. Dashed lines are drawn between pairs of residues with any interresidue distance ≤ 3 Å. Intermolecular pseudobonds are color-coded by PAE value as indicated. (b) Y2H spot assay demonstrating SLY1 and SLY2 interact with SLY1 and SLY2, but not SLXL1 and SLX.

We find SLY1 and SLY2 homodimerize and heterodimerize, but neither interact with SLXL1 or SLX (Fig. 4b), consistent with previous immunoprecipitation experiments, in which SLXL1 and SLX do not pull down SLY1/2 (8, 9). Given their similarity to SLY1/2, we predict that SLXL1 and SLX also dimerize similarly. However, we cannot test this prediction due to autoactivation of BD-SLX/SLXL1 in the Y2H assay (Supp. Fig. 2).

Next, we tested the necessity of the SLY1/2 α-helical domains for dimerization using deleted SLY1/2, which lack their C-terminal α-helices. Deletion of the α-helix of either SLY1 or SLY2 eliminated the interaction with full-length SLY1/2 (Supp. Fig. 7), indicating the necessity of the α-helix domains for binding. SLY1/2 α-helices alone are not sufficient for interaction with full-length SLY1/2, suggesting the N-terminal β-sheet or adjacent unstructured regions are important for dimerization (Supp. Fig. 7).

Because SLXL1, SLX, and SLY1/2 are related to SYCP3, and SLY1 and SLY2 can interact with each other, we tested whether SYCP3 interacts with SLXL1/SLX/SLY1/SLY2 or SPIN1/SSTY1/SSTY2. AlphaFold Multimer predicts SYCP3 interacts with the SYCP3-like proteins (Supp. Fig. 8a) and the Spindlins (Supp. Fig. 8b). However, SYCP3 does not interact with SLXL1/SLX/SLY1/SLY2 or SPIN1/SSTY1/SSTY2 in the Y2H assay (Supp. Fig. 8c). Altogether, our findings suggest the SYCP3-like proteins evolved homo- or hetero multimers to compete.

### SLXL1, SLX, and SLY1/2 proteins compete for Spindlin binding dose-dependently

We have demonstrated specific interactions of SYCP3-like proteins with SPIN1 and SSTY2, which we propose form the basis of competition between the X-linked *Slxl1* and *Slx* genes and the Y-linked *Sly* gene. Next, we tested dose-dependent Spindlin-binding competition between SLXL1/SLX and SLY1/2. While Y3H is typically used to test protein interactions mediated by bridging RNAs or proteins, we modified Y3H as a dosage-regulated competition assay, to test if the expression of a competing protein can disrupt the interaction between two other proteins (Fig. 5a). We have generated a vector (pMFA-Y3H-TetOff) based on pBridge, containing a Tet-Off expression cassette to control expression of a putative competitor gene (Supp. Fig. 9). Expression of the competitor is downregulated in the presence of doxycycline (Dox), allowing us to control the amount of competitor in the system by changing the concentration of Dox. Bait and prey proteins are expressed as in Y2H from pMFA-Y3H-TetOff and pGAD, respectively.

**Figure 5.**
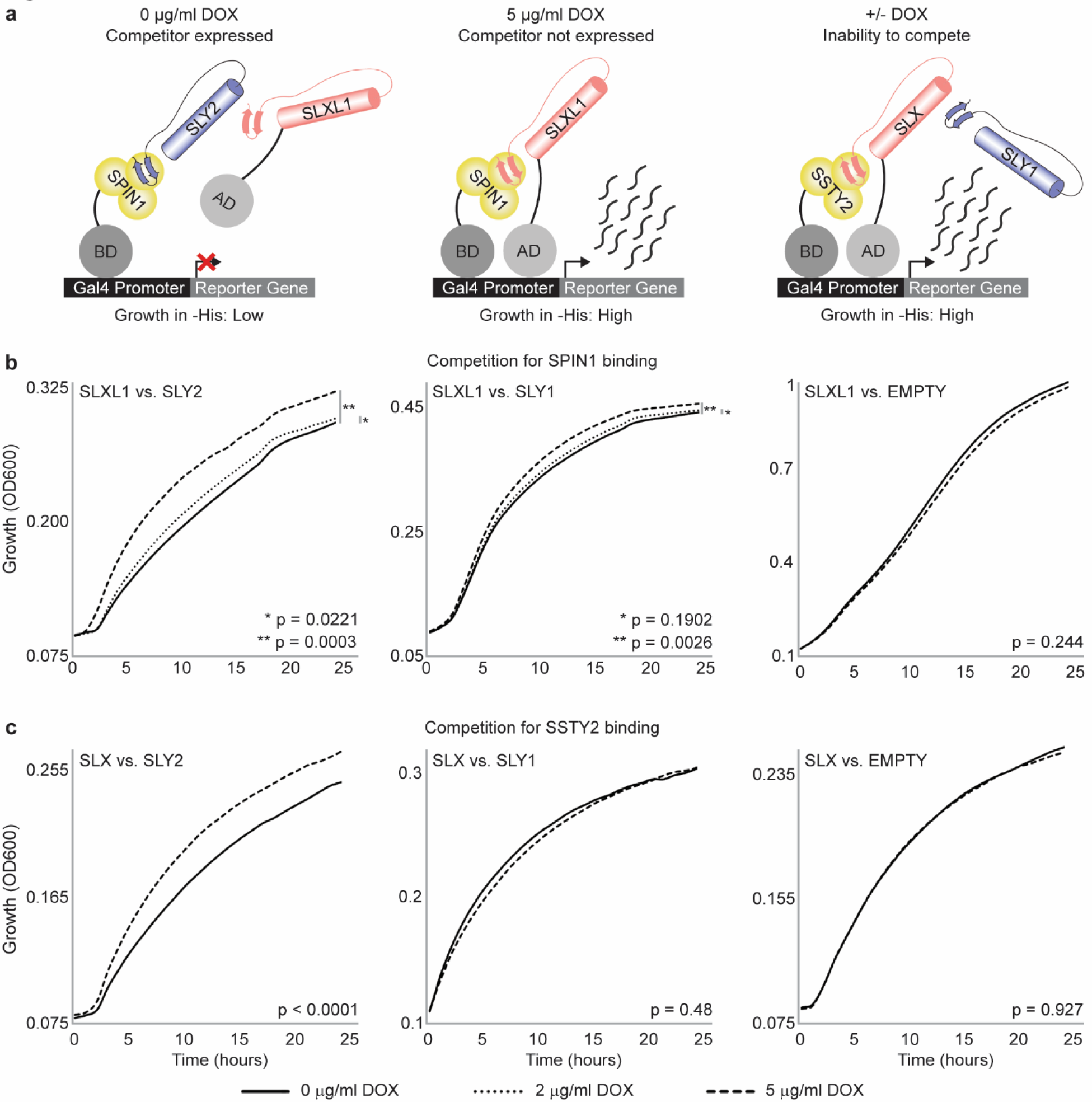
SLX and SLXL1 compete with SLY1/2 for binding to SPIN1 and SSTY2 proteins. (a) Outline of Y3H competition assay. Expression of a competitor protein is controlled by a Tet-Off promoter. When the competitor is expressed (SLY2, left), it blocks interaction of SPIN1 and SLXL1, resulting in reduced *His3* reporter gene expression and low growth. When SLY2 is not expressed (middle), growth rate increases due to better activation of *His3*. If the gene controlled by DOX does not compete (right), the growth rate will remain the same in the presence or absence of DOX. (b) SLY2 (left) and SLY1 (middle) compete with SLXL1 for binding to SPIN1 in a dose-dependent manner, as indicated by increased growth rate with increasing DOX concentration. An EMPTY competitor negative control (right) shows no change in growth rate. (c) SLY2 competes with SLX for binding to SSTY2 (left) in a dose-dependent manner. SLY1 (middle) and EMPTY competitor negative control (right) shows no change in growth rate.

We tested the ability of SLXL1 and SLX to compete with SLY1 and SLY2 for binding SPIN1 and SSTY2. In the case of SLXL1-SPIN1 interactions, expression of SLY1 and SLY2 reduce growth in -His-Ura-Trp media, and Dox increases growth in a dose-dependent manner, consistent with differences in *HIS3* reporter gene expression due to competition (Fig. 5b). Similarly, growth in SLX-SSTY2 interactions is reduced by SLY2, with growth restored by Dox repression of SLY2 expression (Fig. 5c). Importantly, no change in growth is seen when an EMPTY competitor or non-competing gene are used (Fig. 5b,c). These experiments demonstrate that SLY1/2 can compete with SLXL1 for binding to SPIN1, and SLY2 can compete with SLX for binding to SSTY2.

### *Slxl1*, *Slx*, and *Sly* spermatid expressed gene families were recently acquired and amplified

Knowing how SLXL1, SLX, SLY1, and SLY2 proteins interact and compete allows us to test if there are parallels with how each gene family evolved. We first addressed when each gene family arose by building transcriptome assemblies from published testis and sorted testicular germ cell RNA-seq data. We identify only *Slxl1* transcripts in *A. uralensis*, *M. mattheyi*, and *M. pahari*, which map to syntenic regions of *Slxl1* in *M. musculus*. In *M. pahari, Slxl1* transcripts are only detected in sorted spermatocyte-based, not spermatid-based, transcriptome assemblies, indicating *Slxl1* had yet to acquire spermatid-specific expression. In *M. caroli*, we detect transcripts with best reciprocal alignments to *M. musculus Slxl1* and *Sly2*, but not *Sly1* or *Slx*, suggesting the acquisition of *Sly2* in the common ancestor of *M. caroli* and *M. musculus* (Fig. 6a, Supp. Fig. 10a, b). In *M. spretus*, we detect transcripts with best reciprocal matches to *M. musculus Slxl1*, *Slx*, *Sly1,* and *Sly2*, indicating all four gene families were present in the common ancestor of *M. spretus* and *M. musculus*. These findings suggest the competition initiated between SLXL1 and SLY2 in the common ancestor of *M. musculus* and *M. caroli*.

Knowing when each gene family member arose, we used comparative sequence analyses with the *Slxl1* syntenic region of *M*. *caroli*, *M. spretus*, and *M. musculus* to assess how they were acquired. *Sly2* was likely acquired via a transposition of *Slxl1* to the Y chromosome because the intron-exon structure is maintained (Supp. Fig. 10c) (8). *Sly2* gave rise to *Sly1* via an internal duplication of two exons present in *M. musculus* transcripts and genomic DNA but not in *M. caroli* transcripts (Fig. 6b, Supp. Fig. 9c). *Slx* likely arose from *Slxl1*, because of the contiguous genomic alignments of the *M. musculu*s *Slx* coding region with the *M. caroli Slxl1* region and *M. musculu*s *Slx* retains two ancestral *Slxl1* exons, lost in *M. musculus Slxl1* (Supp. Fig. 10b). In the common ancestor of *M. musculus* and *M. spretus,* SLY1 and SLX arose from SLY2 and SLXL1, respectively (Fig. 6b).

**Figure 6.**
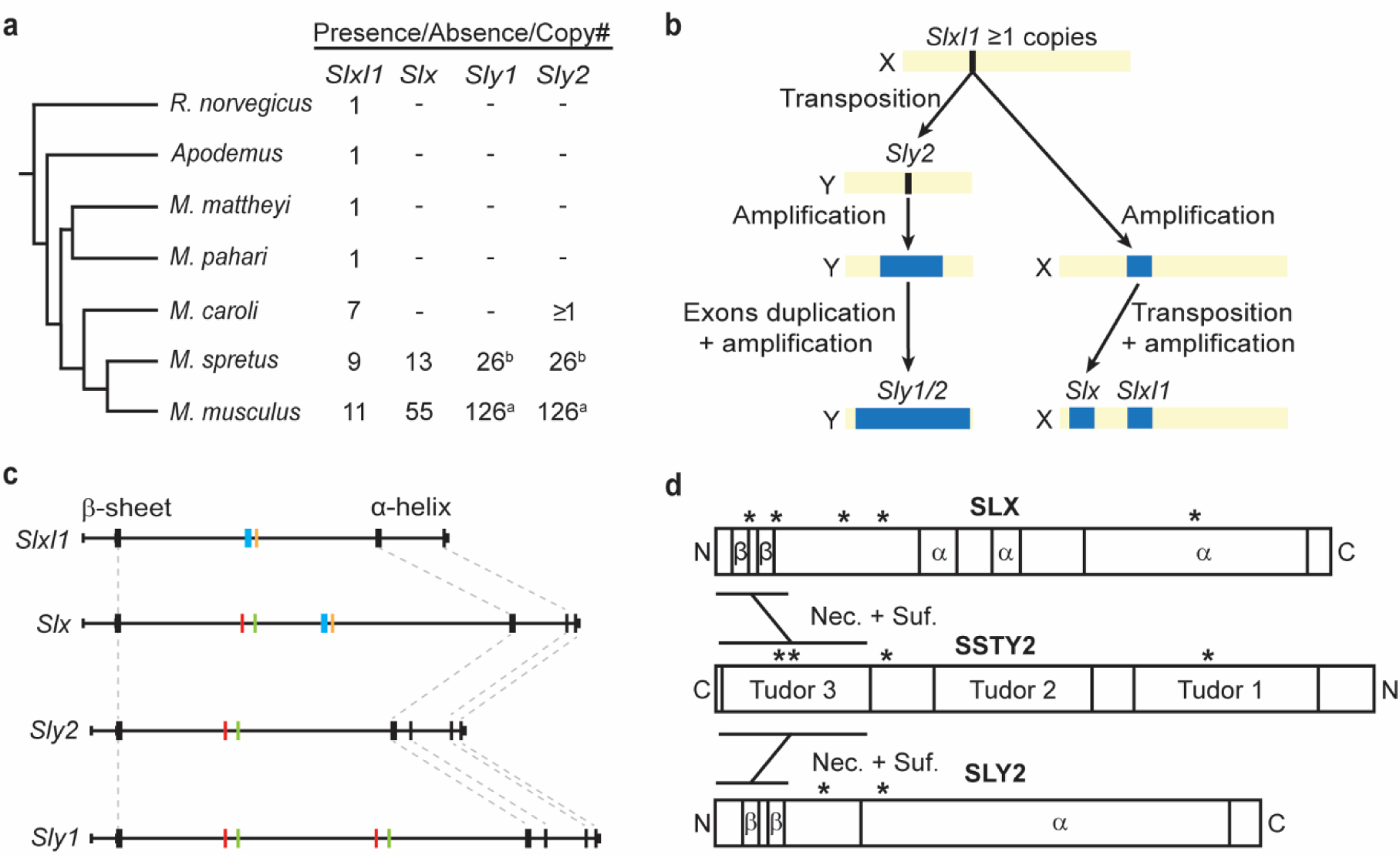
Recent acquisition, amplification, and rapid evolution of *Slx* and *Sly*. (a) *Sycp3*-like gene family members acquisition and amplification history since the divergence with rats, based upon transcriptome comparisons and copy number estimates. *M. musculus* copy number estimates are based on previous studies (11, 43). *M. spretus* and *M. caroli* copy number estimates are based on our read-depth analyses. (b) Sequential events lead to each *Sycp3*-like gene family member’s acquisition. (c) Exon divergence between the gene families occurs outside the β-strands and α-helices. Dotted lines indicate conserved exons, and colored exons indicate duplicated or lost exons. (d) Rapidly evolving residues (* = a >95% posterior probability of being under positive selection) within or near interaction domains. SLX positions are 12D, 20Y, 45L, 59D, and 165Q. SLY2 positions are 38S and 68M. SSTY2 positions are 60G, 168E, 201V, and 205P. (Nec. = necessity; Suf. = sufficiency interaction domains)

We used genome sequence read-depth analyses to support the acquisition events and assess levels of gene amplification. In *M. caroli*, we estimate there are seven copies of *Slxl1* (Fig. 6a). However, there is no increased read depth or sequences matching *Slx* coding sequences in the *M. caroli* regions syntenic to *M. musculus Slx*, consistent with the absence of *Slx* in *M. caroli*. We cannot determine how many copies of *Sly2* are present in *M. caroli* because no Y chromosome sequence is available, but we expect >1 copy because we detect multiple distinct transcripts. In *M. spretus*, *Slxl1*, *Slx*, and *Sly1/2* are all amplified, consistent with previous studies (21). *Sycp3*-like genes amplified upon acquisition, consistent with a dose-dependent competition between X- and Y-linked gene family members (8). While *Slxl1* first evolved around 15 million years ago in the common ancestor of mouse and rat, it recently and rapidly gained spermatid-specific expression and duplicated to evolve *Slx* and *Sly*. Finally, these *Sycp3*-like genes became massively amplified within the last 7 million years (Fig. 6a).

### SLX, SLXL1 and SLY1/2 exon loss/duplication and rapidly evolving residues indicate gene regions under strong selection

Our sequence comparisons revealed the loss and duplication of exons in *Slxl1*, *Slx*, and *Sly1/2*. These exons encode protein sequence between the β-strand and α-helix interaction domains, while the exons encoding the β-strand and α-helix are conserved (Fig. 6c). We suspect the variability in exon usage is driven by the SLX/SLXL1 versus SLY1/2 competition.

SLX/SLXL1 competition with SLY1/2 would suggest that residues under positive selection would be in or near the N-terminal interaction domain and exons that have been gained or lost. We tested which specific residues are under positive selection within each *M. musculus* gene family. The *M. musculus Slx* and *Sly* gene family have a dN/dS of 0.63 and 0.58, respectively, consistent with an elevated dN/dS across the entire gene in cross-species comparisons (8). Most SLX (4/5) and SLY2 (2/2) residues with a >95% posterior probability of being under positive selection (having a dN/dS >1) map within and near the N-terminal interaction domain and within recently gained or lost exons (Fig. 6d). For *Slxl1* there is insufficient variation (only five amino acid differences across the 11 intact *Slxl1* copies) to detect selection. For SSTY2, we observe 3/4 sites under positive selection map within or near Tudor3, the site of interaction with SLX and SLY2 (Fig. 6d). Enrichment of positively selected sites within and near the interaction domains is consistent with N-terminal competition between SLXL1, SLX, and SLY1/2 for the Tudor3 of Spindlins SPIN1 and SSTY2.

## Discussion

This study probes how protein interactions and competition underlie an evolutionary arms race between SLXL1/SLX and SLY1/2 in mice. Understanding the molecular mechanism of this arms race is challenging because the X-linked *Slxl1*, *Slx*, and Y-linked *Sly* gene families are highly duplicated and expressed only in non-culturable haploid spermatids (4, 12, 22). While most previous studies assume SLXL1 and SLX are equivalent in their competition with SLY1/2 (4, 13), we find protein family- and domain-specific molecular interactions. Specifically, SLXL1 competes with SLY1 and SLY2 for binding to SPIN1, and SLX competes with SLY2 for SSTY2 binding. We also discover novel interactions: SLY1/2 can form homo- and hetero-dimers and predict SLXL1/SLX do the same. Based on these data, we propose a model of competing tetramers (Fig. 7a). Since SPIN1 is a chromatin reader (17, 18, 23) and SSTY1/2 are putative chromatin readers (9, 24), we predict these SYCP3-like/Spindlin complexes compete to up- or down-regulate gene expression (Fig. 7B). We have begun to uncover the molecular basis of the competition between SLXL1/SLX and SLY1/2 likely happening in spermatids to influence gene expression and thus X-versus Y-sperm fitness.

**Figure 7.**
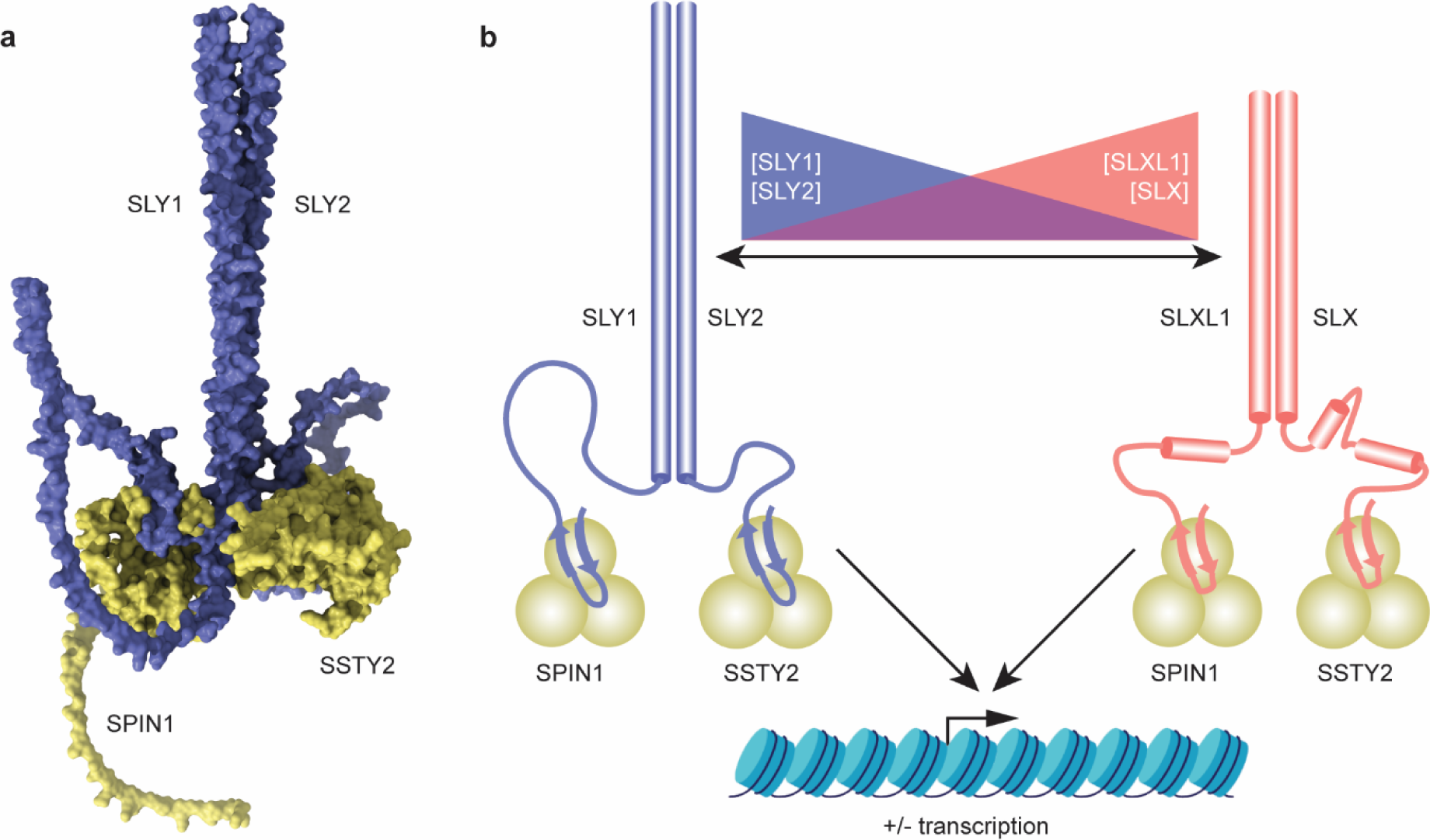
Model of competition between SLXL1/SLX and SLY1/2 for binding to SPIN1/SSTY2. (a) Space-filling model of SLY1/SPIN1 and SLY2/SSTY2 interactions forming a hetero-tetramer via heterodimerization of SLY1 and SLY2, as based on an AlphaFold Multimer predictions. (b) Model illustrating how SLXL1/SLX compete with SLY1/2 for binding to SPIN1/SSTY2. We propose that SLY1-SPIN1, SLY2-SPIN1, and SLY2-SSTY2 can interact as homo-or heterodimers (heterodimers shown here), as can SLXL1-SPIN1, and SLX-SSTY2. However, SLXL1 and SLX cannot dimerize with SLY1/2. The protein complexes containing the X-encoded SLXL1/SLX proteins compete with those containing the Y-encoded SLY1/2 proteins to modulate the ability of Spindlin chromatin readers to interact with specific sites in the genome to potentially up- or down-regulate gene expression.

A striking innovation of the mouse X and Y chromosomes is the high copy number of *Slxl1*, *Slx,* and *Sly* in the evolutionary arms race, suggesting gene dose is an important adaptation/counter-adaptation in this competition (5, 9, 11, 22, 25, 26). We have previously shown that large-scale changes in gene dose impact competition between X- and Y-bearing sperm (8). However, whether the genes or genomic regions are important in dose-dependent competition is unclear. In this study, we demonstrate a dose effect of SLY1/2 when competing with SLXL1/SLX, which helps us understand how gene amplification and dosage contribute to competition at the protein level.

A second potential adaptation in this arms race is the acquisition of sequence variants that increase one protein’s ability to bind its target, making it a better competitor to enhance its transmission. As a counter-adaptation, a competing protein acquires sequence variants that enhance its ability to compete. The high copy number of *Slx*, *Slxl1*, and *Sly* provides many substrates for new mutations, resulting in more competitive alleles. Consistent with sequence variants enhancing binding, the SYCP3-like proteins are rapidly evolving (6, 8), and we find specific residues under strong positive selection are within and near sites of protein interactions. We predict that these sites under strong positive selection will change the binding affinity of SYCP3-like proteins to Spindlins. For example, while SLY1 and SLY2 have a common binding partner in SPIN1, only SLY2 interacts with SSTY2. In our Y3H assay, SLY2 competes better than SLY1 with SLXL1 for SPIN1 binding (Fig. 5b & c). The *Sly* gene family is made up of 28 different sequence variants. We predict different SLY1/2 family members will have improved or diminished competition with SLXL1. Our Y2H/Y3H system allows us to test the effect of these sequence variants on their ability to compete and identify the most competitive alleles. We developed a yeast approach to study competition that can be built upon to deepen our understanding of the mouse *Slxl1/Slx* versus *Sly* arms race and be utilized to study other evolutionary arms races. For example, for *Slxl1/Slx* versus *Sly*, this approach enables us to test the competitive ability of individual alleles and reconstruct the evolutionary arms race in the past ∼3MY. Or we could test whether proteins encoded by *Ssty1/2*, which are in high copy number on the mouse Y chromosome (15, 16), are also in a dose-dependent competition for SPIN1. Beyond the mouse sex chromosomes, this approach can extend to evolutionary arms races across various organisms and systems, particularly in challenging non-model organisms. With only a genome sequence, germline expression data, and cDNA, we can use our Y2H/Y3H approach to identify binding partners of proteins in evolutionary arms races, elucidate the dynamics of the competitive interactions, and understand mechanistically what drives the rapid evolution of their protein sequence. In genomic regions with mazes of mirrored duplicated genes, such as with *Slxl1/Slx* and *Sly*, our yeast-based approach provides a path to understanding how natural selection governs the rapid evolution of these complex genomic regions.

## Materials and Methods

### AlphaFold

AlphaFold Multimer structure predictions of protein interactions were performed using ColabFold-MMseqs2 notebook (27). msa_mode was set to mmseqs2_uniref_env, pair_mode to unpaired_paired, model_type to AlphaFold2-multimer-v2 and num_recycles to 3. AlphaFold outputs were visualized with UCSF ChimeraX (28).

### RT-PCR

Total RNA was extracted from wild-type mouse testes using Trizol (Life Technologies) according to manufacturer’s instructions. Ten mg of total RNA was DNase treated using Turbo DNase (Life Technologies) and reverse transcribed using Superscript II (Invitrogen) using oligo(dT) primers following manufacturer instructions. The coding sequence of each gene was amplified by RT-PCR on adult testis cDNA preparations and cloned into Y2H/Y3H vectors using NEBuilder HiFi DNA Assembly Cloning Kit (New England Biolabs). The specific alleles of each gene family member used in this study are provided in Supp. Table 1. Primers used to clone gene coding sequences are presented in Supp. Table 2.

### Plasmid construction

Bait proteins were expressed from the pBridge vector (Takara), which generates a hybrid protein consisting of the GAL4 DNA-binding domain (BD; amino acids 1–147) and an open reading frame cloned into MCS I. The fusion protein is expressed in yeast host cells from the constitutive ADH1 promoter. The vector also carries the *tRP1* nutritional marker to allow selection of yeast auxotrophs carrying pBridge by growth on limiting synthetic medium lacking tryptophan (-Trp).

Prey proteins were expressed from a modified pGADT7 AD vector (pGAD; Takara), which expresses a protein of interest fused to a GAL4 activation domain (AD; amino acids 768– 881). Transcription of the GAL4 AD fusion is driven by the constitutively active ADH1 promoter. The pGAD vector was modified to replace the *Leu2* selectable marker with *Ura3*, to allow selection of yeast auxotrophs carrying pGAD by growth on limiting synthetic medium lacking uracil (-Ura). All plasmids were sequence verified (GENEWIZ).

### Yeast Two-Hybrid (Y2H)

For Y2H, *S. cerevisiae* AH109 strain was used (*MATa, trp1-901, leu2-3, 112, ura3-52, his3-200, gal4Δ, gal80Δ, LYS2::GAL1UAS -GAL1TATA-HIS3, GAL2UAS-GAL2-TATA -ADE2, URA3:: MEL1UAS-MEL1TATA -lacZ*). Bait and prey plasmids were co-transformed into yeast strain AH109.

Overnight cultures grown in YPDA medium were diluted 1:10 in fresh YPDA and grown until OD600 reached 0.55-0.75. Cells were rinsed twice in 10ml 0.1 M LiAc and resuspended in 0.1 M LiAc and incubated for 10 minutes at room temperature. Approximately 800ng of each plasmid, 100µg of boiled salmon sperm DNA (Invitrogen), and 100µl of cells were mixed. After addition of 700µl of a 0.1 M LiAc/40% PEG3350 mix, cells were incubated for 30 min at 30°C. Cells were mixed with 85µl DMSO and heat shocked at 42°C for 15 minutes and allowed to recover at room temperature. Cells were rinsed with sterile water and plated on selective media.

Growth on -Ura-Trp media selects for both bait and prey plasmids. AH109 cells were co-transformed with pBridge and pGAD expressing different combinations of bait and prey proteins, respectively. To control for autoactivation, one plasmid containing a protein coding gene was mixed with either empty pBridge or empty pGAD. Positive interaction between expressed proteins was assessed by the ability of yeast cells to grow on -His-Ura-Trp (-HUT) and -Ade-His-Ura-Trp (-AHUT).

### Yeast Three-Hybrid (Y3H) competition

We designed a modified pBridge vector (Clontech) in which the PMET25-driven expression cassette was replaced with a Tet-Off expression cassette, which controls the expression of a competitor protein (pMFA-Y3H-TetOff; Supp. Fig. 9; GenBank Accession PQ360726). A tetracycline-controlled transactivator protein (*tTA*), composed of the Tet repressor DNA binding protein (TetR) fused to the VP16 transactivating domain is expressed from a CMV promoter. The tTA regulates the expression of a target gene under transcriptional control of a tetracycline-responsive promoter element (TRE), made up of Tet operator (tetO) sequence concatemers fused to a CYC1 promoter. A coding sequence can be cloned into a NotI site. Expression of the competitor protein is downregulated in the presence of doxycycline (Dox). pMFA-Y3H-TetOff was custom synthesized (Bio Basic) and independently sequence verified (Plasmidsaurus).

Yeast were co-transformed with pMFA-Y3H-TetOff and pGAD plasmids as described above and plated on -HUT plates containing 5 µg/ml Dox. Individual clones were validated by PCR. To generate growth curves, overnight cultures of yeast grown from single colonies in -HUT media to absorbance at 600nm (OD 600) of approximately 0.1 in 200μl of fresh -HUT media with or without Dox and grown in a BioTek Epoch 2 Microplate Spectrophotometer (Agilent) shaking continuously for up to 25 hrs at 30°C. OD600 was measured every hour. Compare Groups of Growth Curves (CGGC) permutation test was used to calculate p-values for pairs of growth curves (29).

### Validation of yeast transformants

After transformation, yeast whole cell extracts were performed on colonies as follows. Individual colonies were picked into 100µl of 0.1% SDS and vortexed at high speed for 20 seconds, incubated at 95°C for 5-10 minutes, and vortexed at high speed for 10 seconds. Samples were centrifuged at high speed. 1µl of the supernatant was used as a template for PCR reactions using primers specific to the plasmids (Supp. Table 2; Supp. Fig. 11).

### Molecular evolutionary analyses

To build transcriptomes, we used publicly available whole testis RNA-seq data for *A. uralensis* (30)(ERP12930), *M. mattheyi* (30) (ERP12930), *M. caroli* (SRP043302) (31) and sorted germ cell RNA-seq for *M. pahari* (32) (SRP323114), and *M. spretus* (32) (SRP323114) as input for *de novo* transcriptome assembly with Trinity (33) using the default parameters. We use tblastN of SLXL1, SLX, SLY1 and SLY2 protein sequences to the de novo transcriptome assemblies to identify orthologous transcripts (Supp. Table 3). Protein alignments were performed using MUSCLE (34).

To test for residues under positive selection in each of the *Slxl1*, *Slx*, and *Sly* gene families in *M. musculus* we used codeml in PAML 4 (35). We aligned all copy members with intact ORF coding sequences for a single gene family and ran models M7 (beta) and M8 (beta & ω) to identify potential residues with a dN/dS >1.

Square dot plots comparing genomic regions were generated using a custom perl script (36).

### *Slx*, *Slxl1* and *Sly1/2* Copy Number Estimates

*M. spretus Slx*, *Slxl1,* and *Sly* and *M. caroli Slx* and *Slxl1* copy numbers were estimated using fastCN (37), a pipeline for copy-number estimation based on read depth. Briefly, to estimate *Slx* and *Slxl1* copy numbers, short-read sequencing data from *M. spretus* (SRA:ERR9880927) and *M. caroli* (SRA:ERR133992) were mapped to their respective female genome assemblies (GCA_921997135.2, GCA_900094665.2) (38, 39) in which RepeatMasker (40), Tandem Repeat Finder (41) and 50-mers with an occurrence > 50 were masked. To estimate *Sly M. spretus* copy number, short-read sequencing data from *M. spretus* (SRA:ERR9880927) was mapped to the *M. musculus* genome reference (mm10), because it contains a Y chromosome assembly while the *M. spretus* assembly does not. Since the *Sly* repeats between *M. spretus* and *M.musculus* are diverged (11), our estimate of *Sly* copy number in *M. spretus* is likely an underestimate. For all copy-number estimates, read depth was normalized for GC content, averaged in windows containing 3kb of unmasked positions, and converted to copy-number estimates based on a defined single-copy sequence dataset. Single-copy control sequences were defined as regions where the read depth was in the full width at half maximum of the read depth distribution for all 3-kb autosomal windows (42).

## Acknowledgements

We would like to acknowledge Isabelle Lamug for technical assistance in generating clones, Lydia Freddolino and Rebecca Hurto for help with generating growth curves, Anuj Kumar for sharing reagents, and David Burke, Allison Cale, Andreas Hochwagen, Erin Jenson, Ann Marie Lawson, John Moran, and Thomas Wilson for helpful comments. This work was supported by National Science Foundation grant 1941796 (J.L.M.), National Institutes of Health grants HD094736 (J.L.M.), HD104339 (C.M.S.), and GM007544 (A.N.K and C.M.S.), and National Science Foundation Graduate Research Fellowship DGE 1256260 (A.N.K.).

## Notes

### Competing Interest Statement

The authors have declared no competing interest.

