## Supplementary Information for "Reenacting a mouse genetic evolutionary arms race in yeast reveals SLXL1/SLX compete with SLY1/2 for binding to Spindlins"

**Supplemental Table 1. Genes cloned into Y2H/Y3H vectors.**

| Vector | Gene | Gencode ID |
|---|---|---|
| pBridge | *Sly1* | ENSMUST00000186004.1[a] |
| | *Sly2* | ENSMUST00000189109.6[a] |
| | *Spin1* | ENSMUST00000095797.5 |
| | *Ssty1* | ENSMUST00000186035.1 |
| | *Ssty2* | ENSMUST00000190565.1 |
| | *Gm4868*[b] | ENSMUST00000164138.3 |
| | *Spin2c* | ENSMUST00000049999.8 |
| | *Spin4* | ENSMUST00000096367.4 |
| | *Sstx* | ENSMUST00000238728.1 |
| pGAD | *Slxl1* | ENSMUST00000088740.4 |
| | *Slx* | ENSMUST00000096913.4 |
| | *Sly1* | ENSMUST00000189109.6[a] |
| | *Sly2* | ENSMUST00000189109.6[a] |
| | *Spin1* | ENSMUST00000095797.5 |
| | *Ssty1* | ENSMUST00000186035.1 |
| | *Ssty2* | ENSMUST00000190565.1 |

[a] ENSMUST00000186004.1 and ENSMUST00000189109.6 have one nucleotide difference that does not change amino acid sequence.
[b] Gm4868 annotated as pseudogene.

**Supplemental Table 2. PCR Primers**

**A. Oligos used to generate protein-coding gene sequences for plasmids**

| Name | Sequence |
|---|---|
| Spin1 BD F | ACAGTTGACTGTATCGCCGGCCATGAAGACCCCATTCGGG |
| Spin1 BD R | CAGGTCGACGGATCCCCGGGTTAGGATGTTTTCACCAAATCGTAG |
| Ssty1 BD F | ACAGTTGACTGTATCGCCGGCCATGTCATCCCTCATGAAG |
| Ssty1 BD R | CAGGTCGACGGATCCCCGGGTTATGATTTTTCAACTTCAAGAATC |
| Ssty2 BD F | ACAGTTGACTGTATCGCCGGCCATGACATCACTCAAGAAG |
| Ssty2 BD R | CAGGTCGACGGATCCCCGGGTTAAGTGATATTTGACACCAG |
| Slxl1 BD F | ACAGTTGACTGTATCGCCGGCCATGGCTCTTAAGAAACTG |
| Slxl1 BD R | CAGGTCGACGGATCCCCGGGTCATTTTCTCAATTCACCATC |
| Slx BD F | ACAGTTGACTGTATCGCCGGCCATGTCTATTAAGAAACTGTGG |
| Slx BD R | CAGGTCGACGGATCCCCGGGTCATAATGTCTCTTCACCATC |
| Sly BD F | ACAGTTGACTGTATCGCCGGCCATGAGAAGAATGGCTCTTAA |
| Sly BD R | CAGGTCGACGGATCCCCGGGTTAGTTCTTGGTCCCCAAGT |
| Spin1 AD F | TATGGCCATGGAGGCCAGTGCCATGAAGACCCCATTCGGG |
| Spin1 AD R | ATCGATGCCCACCCGGGTGGTTAGGATGTTTTCACCAAATCGTAG |
| Ssty1 AD F | TATGGCCATGGAGGCCAGTGCCATGTCATCCCTCATGAAG |
| Ssty1 AD R | ATCGATGCCCACCCGGGTGGTTATGATTTTTCAACTTCAAGAATC |
| Ssty2 AD F | TATGGCCATGGAGGCCAGTGCCATGACATCACTCAAGAAG |
| Ssty2 AD R | ATCGATGCCCACCCGGGTGGTTAAGTGATATTTGACACCAG |
| Slxl1 AD F | TATGGCCATGGAGGCCAGTGCCATGGCTCTTAAGAAACTG |
| Slxl1 AD R | ATCGATGCCCACCCGGGTGGTCATTTTCTCAATTCACCATC |
| Slx AD F | TATGGCCATGGAGGCCAGTGCCATGTCTATTAAGAAACTGTGG |
| Slx AD R | ATCGATGCCCACCCGGGTGGTCATAATGTCTCTTCACCATC |
| Sly AD F | TATGGCCATGGAGGCCAGTGCCATGAGAAGAATGGCTCTTA |
| Sly AD R | ATCGATGCCCACCCGGGTGGTTAGTTCTTGGTCCCCAAG |
| Spin2c BD F | ACAGTTGACTGTATCGCCGGCCATGAAGACCCCTCACAAAAA |
| Spin2c BD R | CAGGTCGACGGATCCCCGGGCTAAGAGTTCTTTTTCACC |
| Spin2E BD F | ACAGTTGACTGTATCGCCGGCCATGGAAAGTTCAAAGCTGA |
| Spin2E BD R | CAGGTCGACGGATCCCCGGGTTAGAACTCTTTCTCCAAGTG |
| Spin4 BD F | ACAGTTGACTGTATCGCCGGCCATGTCTCCTCCGACTGTG |
| Spin4 BD R | CAGGTCGACGGATCCCCGGGTTACGGGGTTTTCACCAAGC |
| Gm4868 BD F | ACAGTTGACTGTATCGCCGGCCATGAAGCCTGAACACAGAATG |
| Gm4868 BD R | CAGGTCGACGGATCCCCGGGTTATTATCCATCTACGTCCCCACAC |
| Slxl1Nterm35 AD R | ATCGATGCCCACCCGGGTGGTCAAGAATGAGCGTCCTCCAAA |
| SlxNterm35 AD R | ATCGATGCCCACCCGGGTGGTCACCTCTTTACTTCAGAATGAGC |
| Sly1Nterm38 AD R | ATCGATGCCCACCCGGGTGGTTACGAAAGAGCCTCCTCTGAAAC |
| Slxl1 Ndel AD F | TATGGCCATGGAGGCCAGTGCCATGGATGATGAAGACGATGACATA |
| Slx Ndel AD F | TATGGCCATGGAGGCCAGTGCCATGGATTCAGATGAAGAAGAAGAG |
| Sly Ndel AD F | TATGGCCATGGAGGCCAGTGCCATGGATGATGAGGACGATGACATA |
| Spin1 Tudor2 BD F | ACAGTTGACTGTATCGCCGGCCATGTCTCGGATCAGCGATGCACA |
| Spin1 Tudor2 BD R | CAGGTCGACGGATCCCCGGGTTAATTGGAATCAGGCATGATGC |
| Spin1 Tudor3 BD F | ACAGTTGACTGTATCGCCGGCCATGGATTCCAATGATTCGCC |
| Ssty2Tudor3 BD F | ACAGTTGACTGTATCGCCGGCCATGATCATTCCAGAGACCC |
| Ssty2delTudor3 BD R | CAGGTCGACGGATCCCCGGGTTAGTGGAGGTTACCTTCCTTG |
| Sly1 Alpha AD F | TATGGCCATGGAGGCCAGTGCCATGTCTGGAGATGACATTTATAAG |
| Sly1 AlphaDel AD R | ATCGATGCCCACCCGGGTGGTTATTTATCCAACATACTGCCTAC |
| Y3H Sly F | TCCTAAGAAGAAGAGAAAGGTGATGAGAAGAATGGCTCTTAA |
| Y3H Sly R | CCGAAGATCTTCGGGCTAATGTTAGTTCTTGGTCCCCAAG |

**B. Oligos used to screen plasmids**

| Name | Sequence |
|---|---|
| pBridge F | CGACATCATCATCGGAAGAG |
| pBridge R | CCTGAGAAAGCAACCTGACC |
| pGAD F | GCCATGGAGTACCCATACGA |
| pGAD R | GCACGATGCACAGTTGAAGT |

| | |
|---|---|
| pBridge Y3H F | GACACGCAAACACAAATACACA |
| pBridge Y3H R | CTTTTCGGTTAGAGCGGATG |
| Slxl1 R2 | CCACTTAACAAATTCCTGTTTTCT |
| Slx R4 | TTGCCAAATGCTGGCCTCTT |
| Sly R2 | ACTTCCGAAAGAGCCTCCTC |
| Spin1 R | GATCCTGCAGCCTACGATGT |
| Ssty1 R | TAAAGAGAAGGGTTTGTTGG |
| Ssty2 R | AGGGTTTGTTGGCAGTTGAC |
| Spin2c R | AATTCTACAGCCCACGATGC |
| Spin2e R | CCAGGTACAGGGAAGGGTTT |
| Spin4 R | GCTCATTGCCTTCTTTCCAG |
| Gm4868 R | GATGGCATGCTCAGCTTCTT |
| Spin1(T3) R | TGGCATATTCCACTTGCTTG |
| Sly1alpha R | TCTTCATTGAATTTCTGTACATCCA |

**Supplemental Table 3:** ***Slxl1/Slx/Sly*** **orthologous transcripts**

```
>Slxl1_Apodemus_uralensis_testis_RNAseq_len=969
CTGTTACCCAGTGTGCTTTGTGGTGGCCACAAGGACGAGTTCTTGACGAAGAAGCCAAGTTCGTGAGGAAGCTTGAA
CATAATCTAGTAATGCATAATAATGGGAATGCAAATGCTGAAGAAATACTTGGAGATACACGATCTCAAGTGCAAAA
TATTGTGGAAAAATTTAAAGGTGACATTAACAAGACTCTTCACGTAAAAAGAAAATCCATGGAAACTTATGTCAAAG
ATTCTTTCAAAGACAGTAATGAAAACTTAGAACAAATTTGGAAAGTGAACAAACGGGAAAGGAAGAAGGTCAACAAC
AAATTTTGTAAGCAGTATATATCTACATTTCAGAAGTTTGATATGGATGTACAGAAATACAATGAAGAACAAACGGG
AAAGGAAGAAGGTCAACAACAAATTTTGTAAGCAGTATATATCTACATTTCAGAAGTTTGATATGGATGTACAGAAA
TACAATGAAGAACAAGAAAATTCAGCTATCAATTTCCAAGCACAACAAAAAACATTTAAACTGTCAACAAGTAGTCA
GAACCAGTCCCTGAAAGCAATTAGAGAATTGCATGAGAGTTTCATGCAGGGTTTGATGAACTTGGACACCAACAACT
ATGATGTGCTTCTTGATGTAGATGGTGAACTGAAAAAGGAAATGTCTGCGTTTAAAAGAAGCATCATGAAGCATACT
CTGAAGTACTCTTCTACTTTTGATACTACTTCAGACTAATGAAAGCATGCATATTTGCACTTGCTAGTGCATTTGTA
ACAGCAAACTAAAAAATTTGTAAATGTTTGAAAGATTCTTTAATCCTTGTTATTCTGATGATTCTAAGAAGGAGGTT
GAACCTGAGTGATGTGATTGCAGTGTTAACTAGGAGGTAGACCTTTAGAATCAAAATAGAAAACCTTTATTATTAAA
CATCTTTAATATTTTTCTTGTTTAAGTGTAAGAACCTCTTTATAT
ORF: 355-732
MYRNTMKNKRERKKVNNKFCKQYISTFQKFDMDVQKYNEEQENSAINFQAQQKTFKLSTSSQNQSLKAIRELHESFM
QGLMNLDTNNYDVLLDVDGELKKEMSAFKRSIMKHTLKYSSTFDTTSD

>Slxl1_Mus_mattheyi_testis_RNAseq_len=1059
TTCCTGCTACCCAAACCCTGTTACCCAGTATGCTCTGTGGTGGCCATGAAGCGAGTTCTTGAGGAAGCCTGAAAACG
TAACTTTCTTAATGGAAGGGCAGCGTTTTCAATATTCTCTAGAGACACCAATGGAAAACTGGAAGTTGTCAAGTGAT
GAAGAGCAAGATGGGAATGCTTCAGAATTGGACCTTTTGGACCTTATTGAAGAACAGAATCCAGTAACTCATGATGA
TGGGAATGCAAATCCTGAAGAAAAAGTTGGAGATACACGATCTCCAGTACAAAATATTCTGGGAAAATTTGAAGGTG
ACATTAACAAGATGCTTCACAAAAAGAGAAAACACTTGGAAACTTATATCAAAGATTCTTTCAAAGGCAGCAACGAA
AAATTAGAACAACTTTGGAAAACGAACAAACGGGAGAGGAAGAAGATCAACAACAAGTTTTTTAAGCAGTATATAAC
TACATTTCAGAAGTTTGATATGGATATACAGAAATTCAATGAAGAACAAGGAAAATCAGAGAACAATTATCAAAAAG
AACAACAAGCATTAAATCTGTCCAAATGTAGTCAGAGCCAGACCCTGAAAAGAATTAAAAATATGCATAAGAAGTCC
ATAAAGGGTTTCATGAAGGTGGAGACCAACAACTACAATATGCTTTTTGATATAGGTGGTGAACTGAGCAAAGAAAT
CTCTGTGTTTAAAAGAGACATCAGGAAGCATACTCTGACGTACGCTTCTGCTTTCCCATTTTCAGACGAATGAAGCA
GCCATATTTTTCACTTGCTGGTATATTTGTAACTGCAAACTAAAAAATTTGCTAACTTTCTTGTCTCCTATGAAACA
GAGTTTGAAAGGCTCTTTAATCCTTGTTATTCTGATGATTCTAAGAAGGAAGATAAACCCCAGTGATGTGATTACAA
AGTTAACTAGGAGGTATACCTTTAGAGTCAAAAGAGAACTCCTTTACTTTGAAACATCTTTAATCATTTTCTTGTTT
AAGTGTAAGAGCCTCTTTATATTAACTTCCAAATAAAATCTAAATGAATAAAAAAAAA
ORF: 89-766
MEGQRFQYSLETPMENWKLSSDEEQDGNASELDLLDLIEEQNPVTHDDGNANPEEKVGDTRSPVQNILGKFEGDINK
MLHKKRKHLETYIKDSFKGSNEKLEQLWKTNKRERKKINNKFFKQYITTFQKFDMDIQKFNEEQGKSENNYQKEQQA
LNLSKCSQSQTLKRIKNMHKKSIKGFMKVETNNYNMLFDIGGELSKEISVFKRDIRKHTLTYASAFPFSDE

>Slxl1_Mus_pahari_spermatocyte_RNAseq_len=955
CAGGTGTCCGTCATGGAAGTGCCCCGCCCATCGGGTCCCTGCCGTTGGCTCCCACGTGAGAACGGGCACGGAGCACG
TGCTAGGGGCGGGGCTACTACGTAATCGGGTGGGCCCATAGCGTTCGGCCAATCAGCAGAGAGCTTGGCCGGGGTCA
GGCTATATTTTCTCCTGCCCAAGGGCCAGGTTTCCTCAGATGCTTCGAGGCTGCGGAGCCGGCGATGTCGACTGCTC
ACCAGAGCTGCTGAGCAAACATCTAAAGATGGTGCCTGGTGGAAGAAAGCATTCTGGGAAATCTGGGAAGCCACCTT
TGGTTGATCAGCCTAAAAAAGCCTTTGACTTTGAGAAAGATGATAAAGATCTGTCTGGTTCAGAAGAAGATGTTGCC
GATGAAAAGGCTCCAGTAATTGAGAAACATGGAAAGAAAAGATCTGCAGGGATAATTGAAGATGTCGGAGGTGAAGT
ACAGAATATGCTGGAAAAATTTGGAGCTGACATCAACAAAGCTCTTCTTGCCAAGAGGAAAAGAATAGAAATGTATA
CCAAAGCTTCTTTCAAAGCCAGTAACCAGAAAATTGAGCAAATTTGGAAAACACAACAAGAGGAAATACAGAAGCTT
AACAATGAATATTCTCAGCAATTTATGAATGTGTTACAGCAATGGGAAGTGGATATACAGAAATTTGAGGAACAAGG
AGAAAAACTATCTAATCTTTTTCGACAACAACAAAAGATTTTTCAGCAGTCTAGAATTGTTCAGAGCCAGAGAATGA
AAGCAATCAAACAGATACATGAACAGTACATAAAGAGTTTGGAAGATGTGGAAAAGAATAATGATAATCTATTTACT
GGCACACAAAGTGAACTTAAAAAAGAAATGGCTATGTTGCAAAAAAAAGTTATGATGGAAACTGTAAGTTCTATTTT
ATATTGCAGAGATGAGGATTAATGGAGGCAC
ORF: 194-946
MLRGCGAGDVDCSPELLSKHLKMVPGGRKHSGKSGKPPLVDQPKKAFDFEKDDKDLSGSEEDVADEKAPVIEKHGKK
RSAGIIEDVGGEVQNMLEKFGADINKALLAKRKRIEMYTKASFKASNQKIEQIWKTQQEEIQKLNNEYSQQFMNVLQ
```

QWEVDIQKFEEQGEKLSNLFRQQQKIFQQSRIVQSQRMKAIKQIHEQYIKSLEDVEKNNDNLFTGTQSELKKEMAML
QKKVMMETVSSILYCRDED

>Sly2_Mus_caroli_testis_RNAseq_len=670
CAAAGGGTCCTTAGAGACCTCCATCCGGGCCTGGCTCCTGACAACGGTTTTTTTCCTTTGAGGATCTGAGCACGGAA
GGGTGCGGTTGGAAGGTGTTCATCTCTTAGATGATCTAGGACTATTAAGTTCTTAAGAGAAGAATGGCGCTTAAGAA
AATGATGGTGATACCAAAGGAAGGTTACTTATTACTTTTGGACTGTGATGATGAAGAAGATGACATAAGTGTTTCAG
AGGAGTCTCTTACTAAAGTAAAGAAGAGCCCAGCATTTGACAAAGATGAGAATATATCGACTCACATAAAAAAGGAT
GGAGATATGCGAGGTGAACTAGACAGTATGTTGGATAAATCTGGAGGTGACATTTATAAGAAGCTTCACATAAAGAA
AAAATGGATGAAAATGTATATGAAAGATTCTTTCAATGGCAGCAACCAGAAATTAGAAAGGCTTTGCAAAAAGAAAA
ATCGAAAGAGGAAGAACATCAATAAAAAAATATGTGATCAGTATATAACTACATTTCAGAAGTCTGATATGGATGTA
TAGAAATTCAATGAAGAAAAAGAAAATCAGCGGGTTTGATGAACTTGGGGACCAACAACTAAGATATGCTTTTTGAT
ACAGATGGTGAACGGAGAAAAAATGTCTGTGTTTAAAAGAGACATCATGGAGAA
ORF: 141-542
MALKKMMVIPKEGYLLLLDCDDEEDDISVSEESLTKVKKSPAFDKDENISTHIKKDGDMRGELDSMLDKSGGDIYKK
LHIKKKWMKMYMKDSFNGSNQKLERLCKKKNRKRKNINKKICDQYITTFQKSDMDV

>Slxl1_Mus_caroli_testis_RNAseq_len=614
CCTCAGAAGCCTCTGTCCAGGCATCTCTGCTGACAACGGTTTTTTACCGTTGAGGAGCTGAGCACGGAAGGGTGCTG
TTGGGAAAGTGTTCACCTCTGAGATAACCTACAACTGAGTGCTTAAGAGAAAAATGGCAAAGAAAATGTGGGTGGTA
CCAAAGAACGGTTACTTATTACTTCTTGACTATGATGAAGATGAAGATGACATAAATGTTTTGGATGAGGCTCATTC
TGAAGTACAGAATCCAGTAACTCATGATGATGGGAATGCAAATCCTGAAGGAATAGTTGGAGATACACGTGAGATGA
TCAACAACAAATTATGTGAGCAGTATATAACTACATTTCAGAAGTCTGATATGGATATACAGAAATTCAATGAAGAA
CAAGAAAAATCAGTGGGTTTGATGAACTTGGATACCAACAACTATGATAAGCTTTTTGATGTAGATGGTGAACAGAG
AAAATAAATATCTGTGTTTAAAAGAGACATTATGAAGCATACTCTGAAGTACTCATCTACTTTCCCATCTTCAGACT
AATGAAGCATACATATTTTTTACTTGCTGGTACATTTGTAACCAAATAAAAAAATTTGCTAACTTTTTTTCCCCA
ORF: 131-469
MAKKMWVVPKNGYLLLLDYDEDEDDINVLDEAHSEVQNPVTHDDGNANPEGIVGDTREMINNKLCEQYITTFQKSDM
DIQKFNEEQEKSVGLMNLDTNNYDKLFDVDGEQRK

>Sly1_Mus_spretus_spermatid_RNAseq_len=1183
CCTTTACCCGCTTACGGGATGAACCGGTTTTTTTTTTTTTTTTTTTTTTGCCATTGAGGAGCTAAGGACAGAAGGGT
GCGGTTGGGAAGGTGTTCTCCTCTTAGATGAGCTAAGACTACTGAGTTCTTAAGAGAAGAATGGCACTTAAGAAATT
GAAGGTGATACCAAAGGAAGGCTACTTATTACTTTTGGACTTTGATTATGAGGATTACATAAAAGTTTCAGAGGAGG
CTCTATCGGAAGTAAAGAGCCCAGCATTTGATAAAAATGAGAATATATCGCCTCAAGCAGAAGCAGATGAAGATATG
GGGAGGAGGTCCCCTCGAGATCCTCGAATAACTGGACCTGCTGGACGGGTCAAGTGGCGCCCGACGTGGAGCACGAG
GTACGACTGCCCTCCGGACAACGGATTAAGGATTAAAGAGGTACCGCGCAGACAGAGAAACAGTGGGACAGTCCACC
GACACTTCGCTCGAGTGTCGAGTACGGATGTCGGGAGAGATGAAGTAGACAGTATGTTGGATAAATCTGGAGTAAAG
AGCCCAGCACTTGGTAAAGATGAGAATATATCGCCTCAAGTAAAAGGAGATGAAGACATGGGACATGAAGTAGGCAG
TATGTTGGATAAATCTGGAGATGACATTTATAAGACGCTTCACATAAAGAGAAAATGGATGGAAAGTTATATCAAAG
AATCTTTCAAAGGCAGCAACCAGAAATTAGAAAGATTTTGCAAAATGAACAAACGAGAGAGGAAGAACATCAACAAC
AAATTTTGTGAGCAGTATATAACTACATTTCAGAAGTCTGATATGGATGTACAGAAGTTCAATGAAGAAAAAGAAAA
ATCAGTGAATAATTTTCAAAAAGAGCATGCATTGAAACTGTCCAAATGCAGTCAGAACCAGACCCTGGAAGCAGTTA
AAGATATGCATGAGAAGTCCATGGAGGGTTTGATGAACTTGGGGACCAAGAACTAAAATATGCTTTTTGGTGTAGAC
GGTGAACTGAGAAAAAAAAATGTCTATGTTTAAAAGAGCCATCATGGAGAATACTCTGAAGTACTCTTCTACTTTCC
CATCTTCAGAAAAATGAAGCATGAAAATTTTTCACTTGCTGGTATATATATAAAACAAATAAAAAAAAAATCTCTAA
CTTTTTTGTCTCTTATGAAAAAAAAAAA
ORF: 138-980
MALKKLKVIPKEGYLLLLDFDYEDYIKVSEEALSEVKSPAFDKNENISPQAEADEDMGRRSPRDPRITGPAGRVKWR
PTWSTRYDCPPDNGLRIKEVPRRQRNSGTVHRHFARVSSTDVGRDEVDSMLDKSGVKSPALGKDENISPQVKGDEDM
GHEVGSMLDKSGDDIYKTLHIKRKWMESYIKESFKGSNQKLERFCKMNKRERKNINNKFCEQYITTFQKSDMDVQKF
NEEKEKSVNNFQKEHALKLSKCSQNQTLEAVKDMHEKSMEGLMNLGTKN

>Sly2_Mus_spretus_spermatid_RNAseq_len=866
CCTTTACCCGCTTACGGGATGAACCGGTTTTTTTTTTTTTTTTTTTTTTGCCATTGAGGAGCTAAGGACAGAAGGGT
GCGGTTGGGAAGGTGTTCTCCTCTTAGATGAGCTAAGACTACTGAGTTCTTAAGAGAAGAATGGCACTTAAGAAATT
GAAGGTGATACCAAAGGAAGGCTACTTATTACTTTTGGACTTTGATTATGAGGATTACATAAAAGTTTCAGAGGAGG

CTCTATCGGAAGTAAAGAGCCCAGCATTTGATAAAAATGAGAATATATCGCCTCAAGCAGAAGCAGATGAAGATATG
GGGAGGAGGTCCCCTCGAGATCCTCGAATAACTGGACCTGCTGGACGGGTCAAGTGGCGCCCGACGTGGAGCACGAG
GTACGACTGCCCTCCGGACAACGGATTAAGGATTAAAGAGGTACCGCGCAGACAGAGAAACAGTGGGACAGTCCACC
GACACTTCACTCGAGTGTCGAGTACGGACGTCGGGAGAGATGAAGTAGACAGTATGTTGGATAAATCTGGAGATGAC
ATTTATAAGACGCTTCACATAAAGAGAAAATGGATGGAAAGTTATATCAAAGAATCTTTCAAAGGCAGCAACCAGAA
ATTAGAAAGATTTTGCAAAATGAACAAACGAGAGAGGAAGAACATCAACAACAAATTTTGTGAGCAGTATATAACTA
CATTTCAGAAGTCTGATATGGATGTACAGAAGTTCAATGAAGAAAAAGAAAAATCAGTGAATAATTTTCAAAAAGAG
CATGCATTGAAACTGTCCAAATGCAGTCAGAACCAGACCCTGGAAGCAGTTAAAGATATGCATGAAAAGTCCATGGA
GGGTTTGATGAATTGGGGA
ORF: 138-866
MALKKLKVIPKEGYLLLLDFDYEDYIKVSEEALSEVKSPAFDKNENISPQAEADEDMGRRSPRDPRITGPAGRVKWR
PTWSTRYDCPPDNGLRIKEVPRRQRNSGTVHRHFTRVSSTDVGRDEVDSMLDKSGDDIYKTLHIKRKWMESYIKESF
KGSNQKLERFCKMNKRERKNINNKFCEQYITTFQKSDMDVQKFNEEKEKSVNNFQKEHALKLSKCSQNQTLEAVKDM
HEKSMEGLMNWG

>Slxl1_Mus_spretus_spermatid_RNAseq_len=568
>TRINITY_DN4835_c0_g1_i19 len=568 M. spretus Slxl1 (spermatid RNA-seq)
AAATGAAGATATAAGAGATGAACAAGACAGTATGTTGGATAAATCTGGAGAAAACGTAAGTTTCTCAGTGGAATGGC
AGCGTTTTGCAAGTTCTGTAGAGACACCAATAGAAAACAGGAATTTGTTAAGTGGTGAACAGCAAGTTAGGAATGCT
TCAGAATTGGACCTTATGGAAGTACAGAATTCAGTAACTCATGATGATGAGAATGAAATTCCTGAAGAAATAGTTGG
AGATACACGGGAGATGATCAACAGCAAGTCGTGTGAGCAGAATAAAACTACATTTCAGAAGTTTGATATGGATGTCC
AGAATTTCAATGAACAGCAAGAAAAAATCAGTGAGTTTGATGAACTTGGAGACCAACATCGACGATATGCTTTTTGAT
GTAGATGATGAATTGAGAAAATGAATGTCTGTGTTTAAAAGAGACATCATGAAGCATACTCTGAAGTTCTCTTCTCC
TTTCCCATCTTCAGACTAATGAAGCATGCAAATTTTTCACTTGATGGTACATTTGTAAGCAAATAAAAAATTCGCTA
ACTTTTTGTCTCTTATGAAAAAAAAAAAA
ORF: 32-409
MLDKSGENVSFSVEWQRFASSVETPIENRNLLSGEQQVRNASELDLMEVQNSVTHDDENEIPEEIVGDTREMINSKS
CEQNKTTFQKFDMDVQNFNEQQEKSVSLMNLETNIDDMLFDVDDELRK

>Slx_Mus_spretus_spermatid_RNAseq_len=649
AAATGAAGATATAAGAGATGAACAAGACAGTATGTTGGATAAATCTGGAGAAAACGTAAGTTTCTCAGTGGAATGGC
AGCGTTTTGCACGTTCTGTAGAGACACCAATGGAAAACTGGAATTTGTTAAGTGGTGAACAGCAAGTTAGGAATGCT
TCAGAATTGGACCTTATGGAAGTACAGAATTCAGTAACTCATGATGATGGGAATGCAAATCCTGAAGAAGTAGTTGG
AGATACACGAAAGAAGATCAACAACAAATTGTGTGAGCAGAAGTTTGATATGGATATACAGAAATTCAATGAAGAAC
AAGAAAAAATCAGTGAACAATTATCAAAAAGAACAACAAGCATTAAAACTTTCTGAATGTAGTCAGAGTCAGACCCTG
GAAGCAATTGAAGACATGCATGAGAAGTCCATGCAGGGTTTGATGAACATGGAGACCAACAACTACGATATGCTTTT
TGATGTAGATGATGAATTGAGAAAATGAATGTCTGTGTTTAAAAGAGACATCATGAAGCATACTCTGAAGTTCTCTT
CTCCTTTCCCATCTTCAGACTAATGAAGCATGCAAATTTTTCACTTGATGGTACATTTGTAAGCAAATAAAAAATTC
GCTAACTTTTTGTCTCTTATGAAAAAAAAAAAA
ORF: 32-490
MLDKSGENVSFSVEWQRFARSVETPMENWNLLSGEQQVRNASELDLMEVQNSVTHDDGNANPEEVVGDTRKKINNKL
CEQKFDMDIQKFNEEQEKSVNNYQKEQQALKLSECSQSQTLEAIEDMHEKSMQGLMNMETNNYDMLFDVDDELRK

### Supplemental Figures

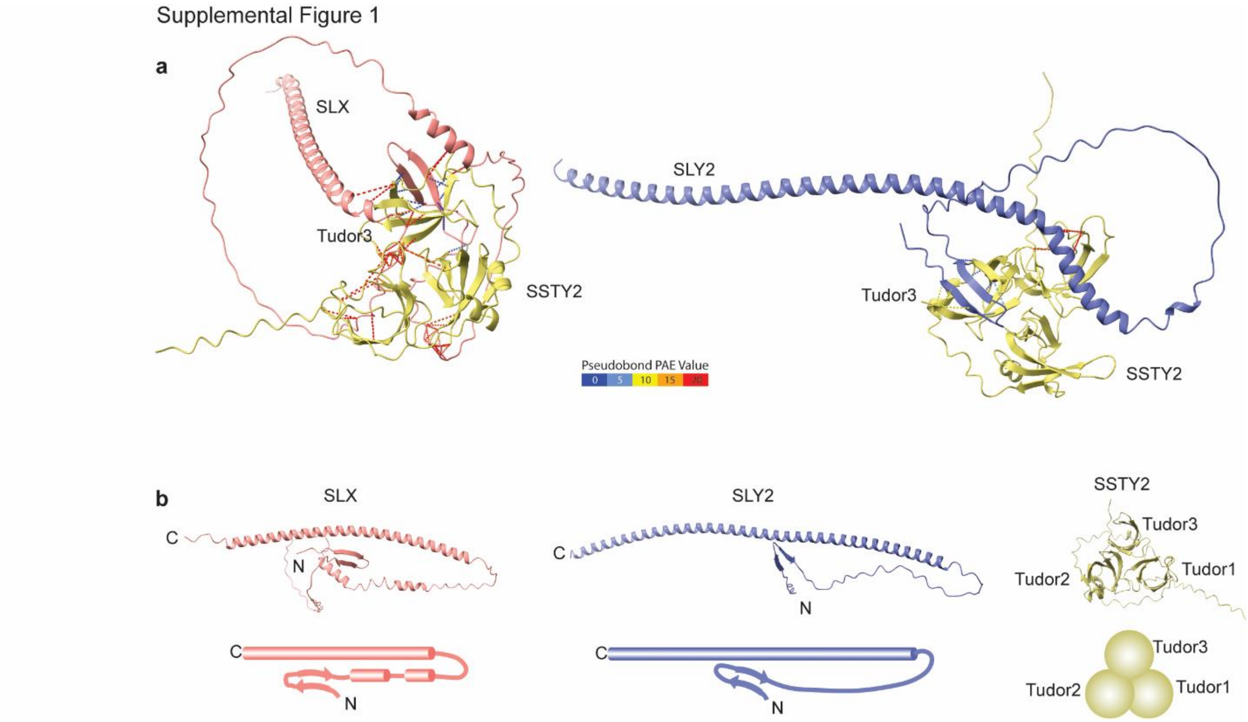


**Supplemental Figure 1. AlphaFold Multimer predicts interactions between SLX/SLY2 and SSTY2.**

(a) AlphaFold Multimer predicts binding of SSTY2 to SLX (left) and SLY2 (right). (b) AlphaFold predictions and cartoons of the 3D structures of SLX, SLY2, and SSTY2.

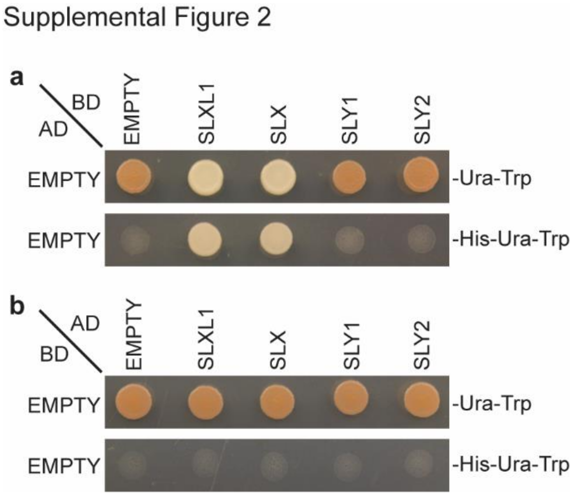


**Supplemental Figure 2. SLXL1 and SLX autoactivate *His3* when linked to Gal4-BD, but not Gal4-AD.**

Y2H spot assays testing if SLXL1, SLX, SLY1, and SLY2 autoactivate the *His3* reporter. (a) SLXL1/SLX/SLY1/SLY2 linked to Gal4-BD co-transformed with empty Gal4-AD vector into yeast strain AH109. SLXL1 and SLX, but not SLY1 and SLY2, can grow without His, indicating SLXL1 and SLX autoactivate with the Gal4-BD. (b) SLXL1/SLX/SLY1/SLY2 linked to Gal4-AD co-transformed with empty Gal4-BD vector into yeast strain AH109. SLXL1, SLX, SLY1, and SLY2 do not grow without His, indicating SLXL1 and SLX do not autoactivate with the Gal4-AD.

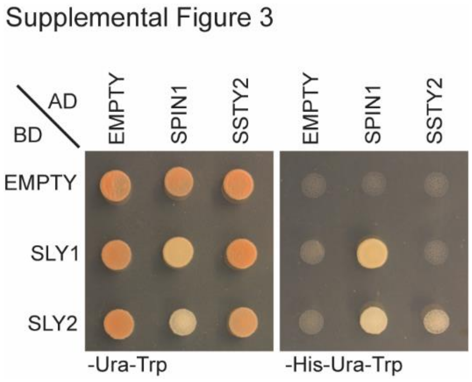


**Supplemental Figure 3. SPIN1 interactions with SLY1/2 and SSTY2 interactions with SLY2 are maintained when BD and AD vectors are swapped.**

Y2H spot assay demonstrating growth on -His-Ura-Trp media indicating an interaction between SPIN1 and SLY1/2 and SSTY2 with SLY2 when BD and AD vectors are swapped. SPIN1 and SSTY2 are linked to the Gal4-AD, and SLY1/2 are linked to the Gal4-BD.

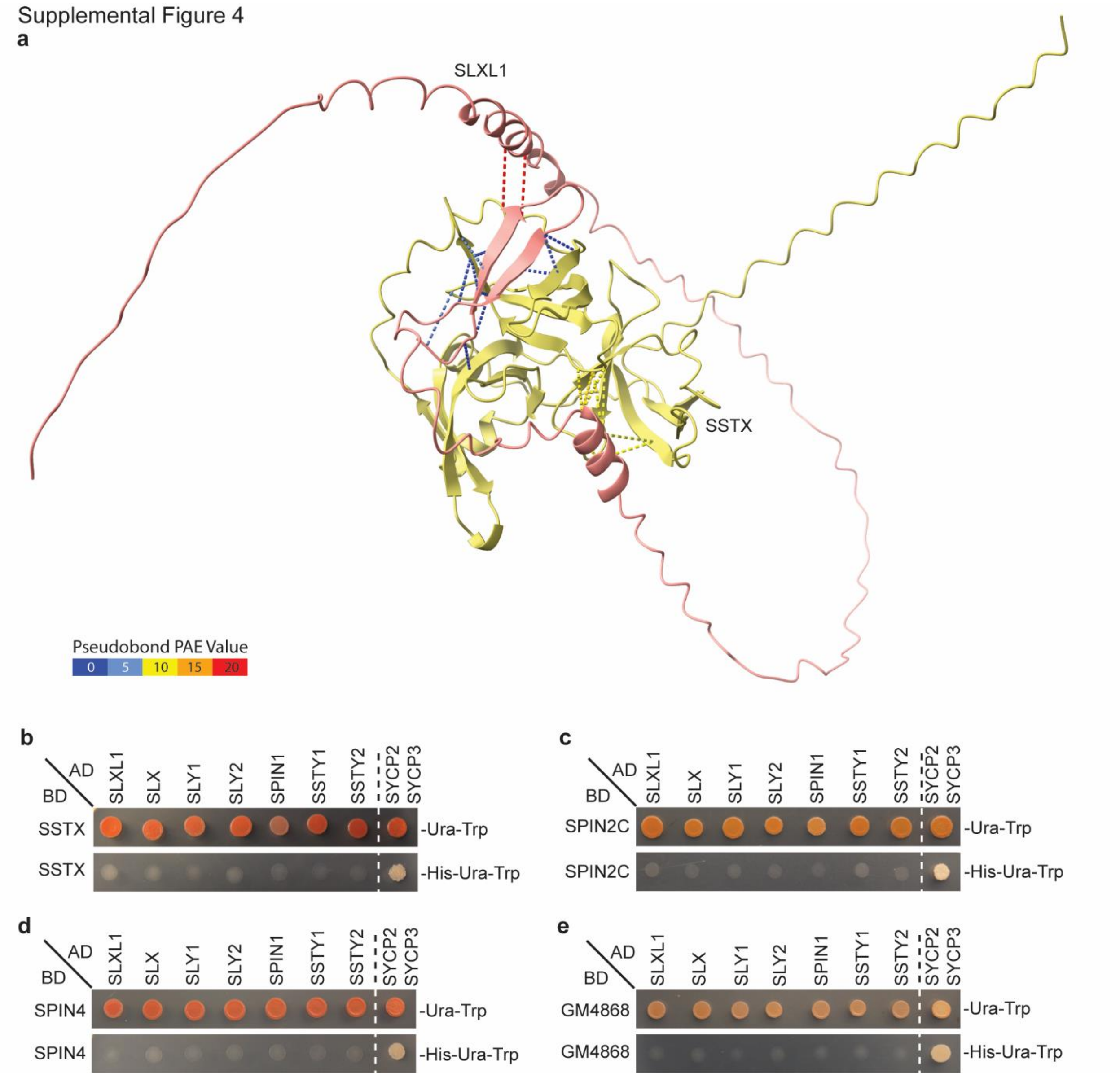


**Supplemental Figure 4. SLXL1/SLX/SLY1/SLY2 do not interact with other Spindlins.**

(a) AlphaFold Multimer prediction of binding of SLXL1 to SSTX. Intermolecular pseudobonds are represented by dashed lines drawn between pairs of residues with any interresidue distance ≤ 3 Å and colored based on pseudobond PAE value. (b-e) Y2H spot assays testing if SLXL1/SLX/SLY1/SLY2 interacts with other Spindlins: (b) SSTX; (c) SPIN2C; (d) SPIN4; (e)

GM4868. Growth on -Ura-Trp indicates the presence of BD and AD vectors. Growth on -His-Ura-Trp indicates binding between BD- and AD-linked proteins. Positive control: SYCP3-BD, SYCP2-AD. Negative controls: Empty vectors.

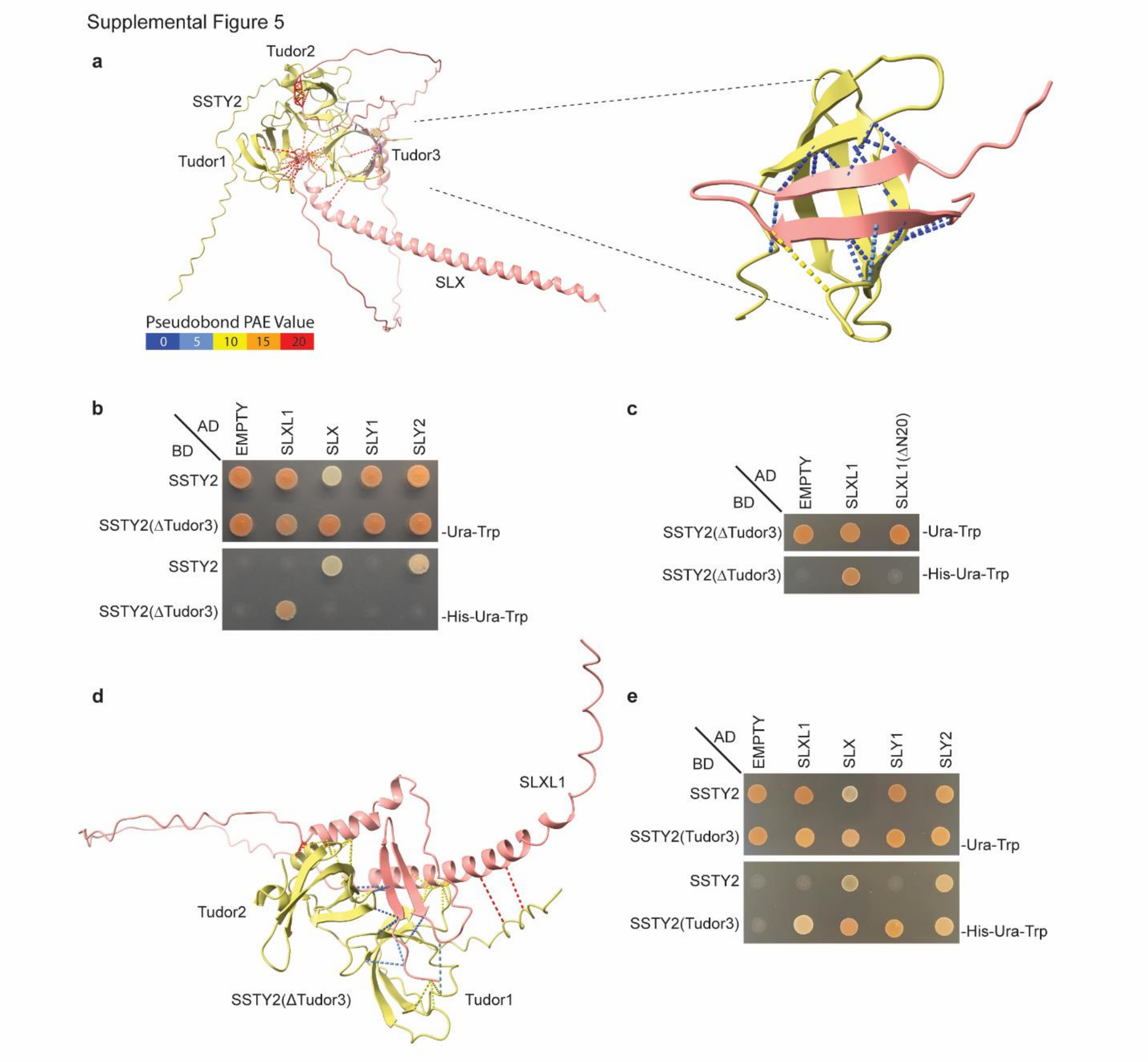


**Supplemental Figure 5. SSTY2 Tudor3 domain is necessary and sufficient for binding to SLXL1/SLX/SLY1/SLY2.**

(a) Predicted interactions between SLX and SSTY2 (left) with a close-up showing interactions between SLXL1 N-terminal β-strands and SPIN1 Tudor3 (right), determined with AlphaFold Multimer. Dashed lines indicate interresidue distance ≤ 3 Å. Intermolecular pseudobonds are color-coded by predicted aligned error (PAE) value as indicated. (b) Y2H spot assays demonstrate SSTY2 Tudor3 is necessary to bind SLX/SLY1/SLY2. SSTY2 lacking Tudor3 can bind SLXL1. (c) Y2H spot assays demonstrate the N-terminal 20 amino acids of SLXL1 are necessary for SSTY2(ΔTudor3) binding. (d) Predicted interactions between SSTY2(ΔTudor3) and SLXL1, mediated by the N-terminal 20 amino acids of SLXL1, determined with AlphaFold Multimer. (e) Y2H spot assays demonstrate SSTY2 Tudor3 is sufficient to interact with SLXL1, SLX, and SLY1/2.

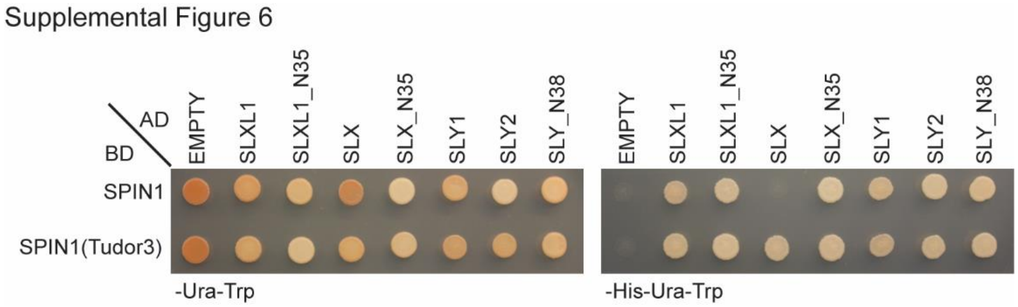


**Supplemental Figure 6. SPIN1 Tudor3 is sufficient to bind the N-termini of SLXL1, SLX, and SLY1/2.**

Y2H spot assay testing if SPIN1(Tudor3) is sufficient to interact with amino acids 1-35 of SLXL1 and SLX or amino acids 1-38 of SLY1/2. SPIN1(Tudor3) can bind to the N-terminal peptides of SLXL1, SLX, and SLY1/2.

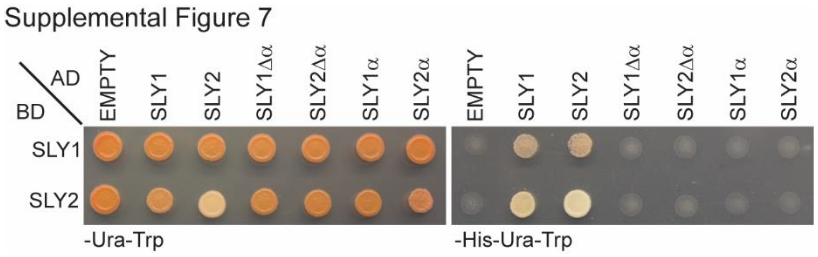


**Supplemental Figure 7. SLY1/2 α-helical domains are necessary but not sufficient for homo- and heterodimer formation.**

Y2H spot assay testing if full-length SLY1 and SLY2 interact with SLY1 and SLY2 lacking the entire α-helix domain (SLY1Δα and SLY2Δα) or with SLY1 and SLY2 α-helix domains alone (SLY1α and SLY2α). SLY1 and SLY2 do not interact with SLY1Δα, SLY2Δα, SLY1α, or SLY2α.

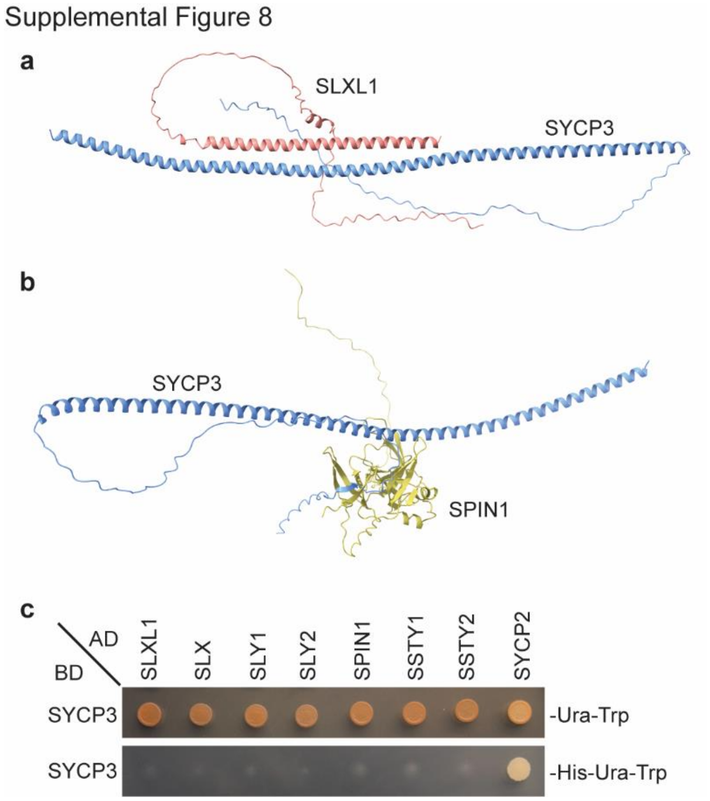


**Supplemental Figure 8. SYCP3 does not interact with SYCP3-like proteins or Spindlins.**

Predicted interactions of SYCP3 with (a) SLXL1 and (b) SPIN1 using AlphaFold Multimer. Intermolecular pseudobonds are represented by dashed lines indicate interresidue distance ≤ 3 Å and colored based on pseudobond PAE value. (c) SYCP3 does not interact with SYCP3-like proteins (SLXL1, SLX, SLY1/2) and Spindlins (SPIN1, SSTY1/2) in Y2H. Positive control: SYCP3-BD, SYCP2-AD. Negative controls: Empty vectors.

Supplemental Figure 9

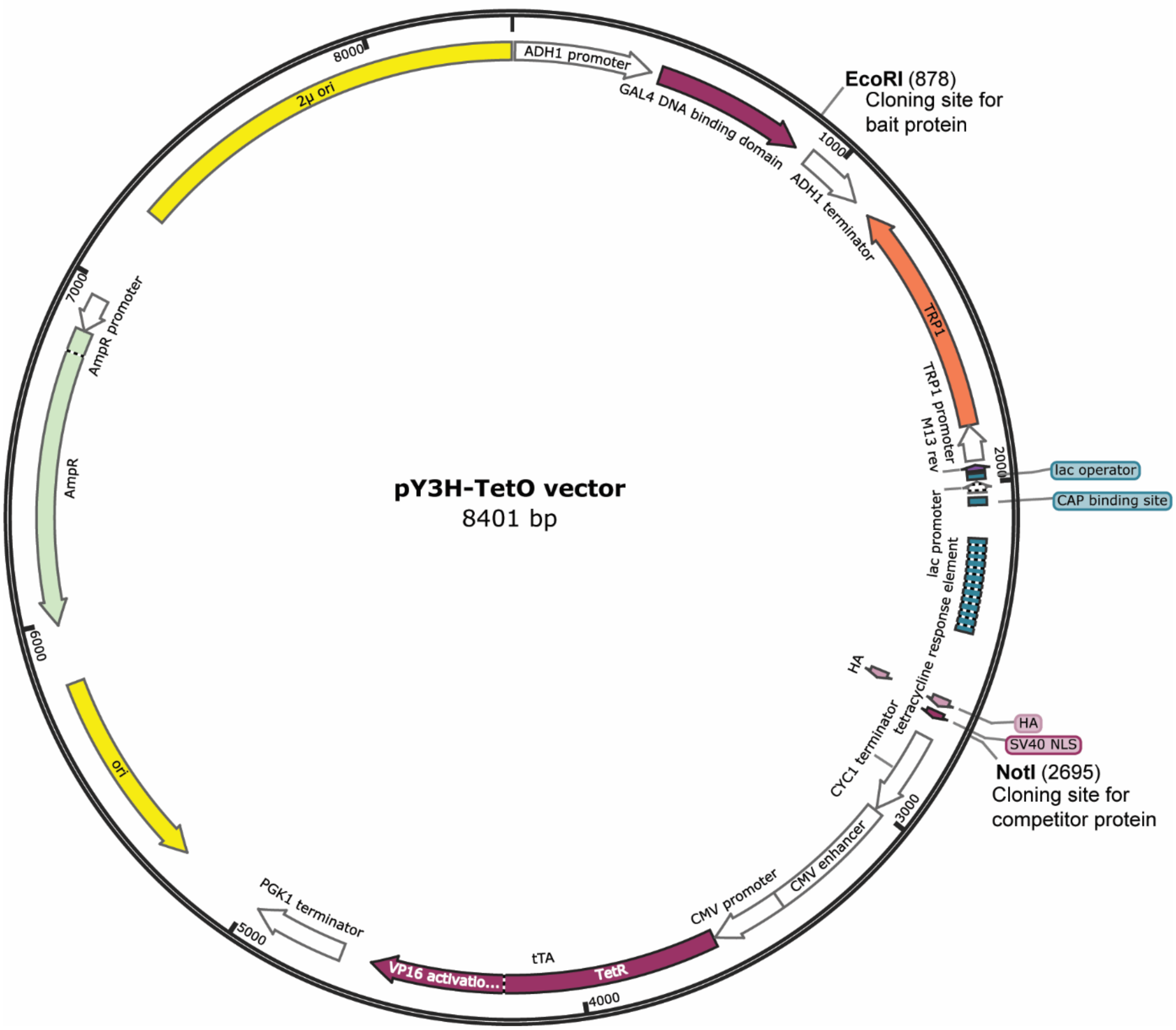


**Supplemental Figure 9. Plasmid map of pBridge Y3H vector.**

Map of the pMFA-Y3H-TetOff vector used for Y3H assay. This vector is modified from the pBridge vector (Clontech). A coding sequence encoding the bait is cloned into the EcoRI site and fused to the cDNA encoding the GAL4 DNA binding domain (DNA-BD). Its expression is driven by the constitutive ADH1 promoter. The vector carries the ampicillin resistance gene (AmpR) for selection in *E. coli*, and the TRP1 nutritional marker for selection in yeast. A second coding sequence for a competitor protein is cloned into a NotI site, where it is fused to a nuclear localization signal and a HA tag. Expression of the competitor is controlled by a Tet-Off system. A tetracycline-controlled transactivator protein (*tTA*), which is composed of the Tet repressor

DNA binding protein (TetR) from the Tc resistance operon of *Escherichia coli* transposon Tn10 fused to the strong transactivating domain of VP16 from Herpes simplex virus, regulates expression of the competitor that is under transcriptional control of a tetracycline-responsive promoter element.

Supplemental Figure 10

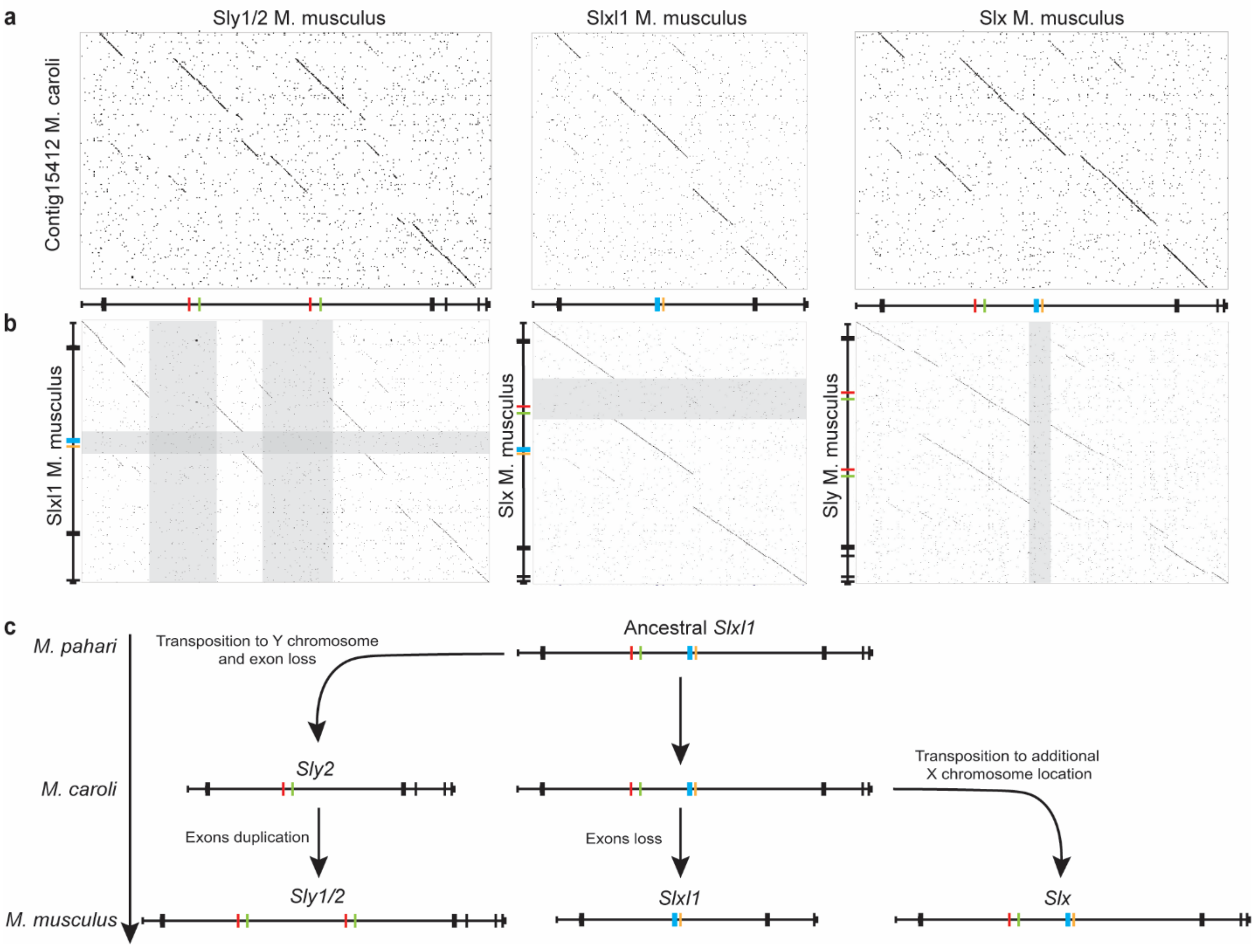


**Supplemental Figure 10. *Slxl1*, *Slx*, and *Sly* lost and duplicated exons during their recent evolution.**

(a) Square plots (where each dot represents 10bp of perfect nucleotide identity representing a dot) of genomic regions spanning *M. musculus Slxl1*, *Slx*, and *Sly* compared to a *M. caroli* contig containing *Slxl1* sequence. (b) Square plots (with 10bp of perfect nucleotide identity representing a dot) of genomic regions spanning *M. musculus Slxl1*, *Slx*, and *Sly* compared to themselves. Genomic regions where exons are lost are shown shaded. (c) Gene structure evolution of *Slxl1*, *Slx*, and *Sly* based on the square plots from panels a and b.

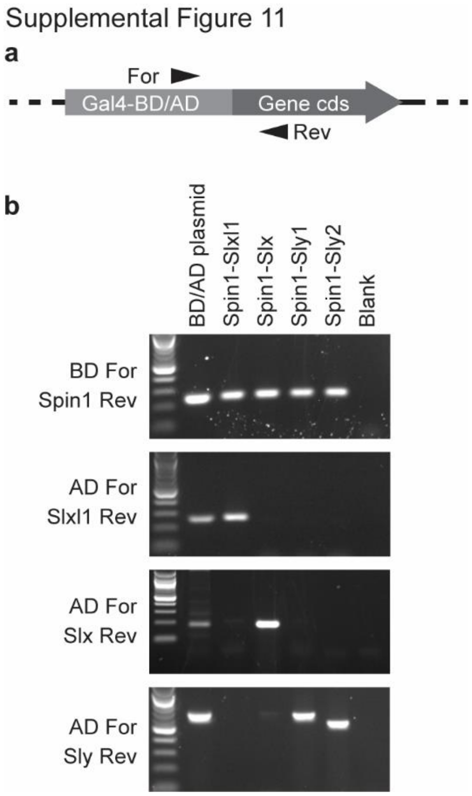


**Supplemental Figure 11. Representative PCR assay confirming successful transformation of Y2H plasmids into yeast.**

(a) Generalized diagram of a gene cds cloned into a Y2H plasmid. Arrows indicate the position of a forward (For) primer located in either the Gal4-Binding Domain (Gal4-BD; Bait) or the Gal4-Activation Domain (Gal4-AD; Prey) and reverse (Rev) PCR primer located within the gene cds. (b) Representative bands from PCR products generated by colony PCR of yeast transformed with pBridge-SPIN1 plus one of the following prey plasmids: pGAD-SLXL1, pGAD-SLX, pGAD-SLY1, pGAD-SLY2. Positive control: purified plasmid DNA. Negative control: water.
